# Long-lived adult-born hippocampal neurons promote successful cognitive aging

**DOI:** 10.1101/2024.07.19.604332

**Authors:** Nicolas Blin, Vanessa Charrier, Fanny Farrugia, Estelle Cartier, Emilie Pacary, Muriel Koehl, Carlos Lois, Dieter Chichung Lie, Nuria Masachs, Djoher Nora Abrous

**Author notes:** Electronic address.

## Abstract

Aging is commonly associated with a decline in memory abilities, yet some individuals remain resilient with preserved memory abilities. Memory processing is critically dependent on adult neurogenesis, a unique form of plasticity in the hippocampus. However, it remains unknown if cognitive aging influences the integration and role of adult-born hippocampal neurons (ABNs) generated early in adult life. Here, we investigated the role of long-lived ABNs in rats characterized as either resilient or vulnerable to cognitive aging using a peudo-longitudinal approach. Our findings reveal that long-lived ABNs support successful cognitive aging by preserving their synaptic inputs onto the proximal segments of their dendrites, and that these proximal synaptic sites also demonstrate a maintenance of their mitochondrial homeostasis. Furthermore, by-passing the reduced inputs of ABNs in vulnerable rats through direct optogenetic stimulation successfully improved their memory abilities. Overall, our data indicate that the maintenance of long-lived ABNs integration within the neuronal network is essential for successful cognitive aging, highlighting their potential as a therapeutic target for restoring cognitive functions in old age.

**Graphical abstract:** 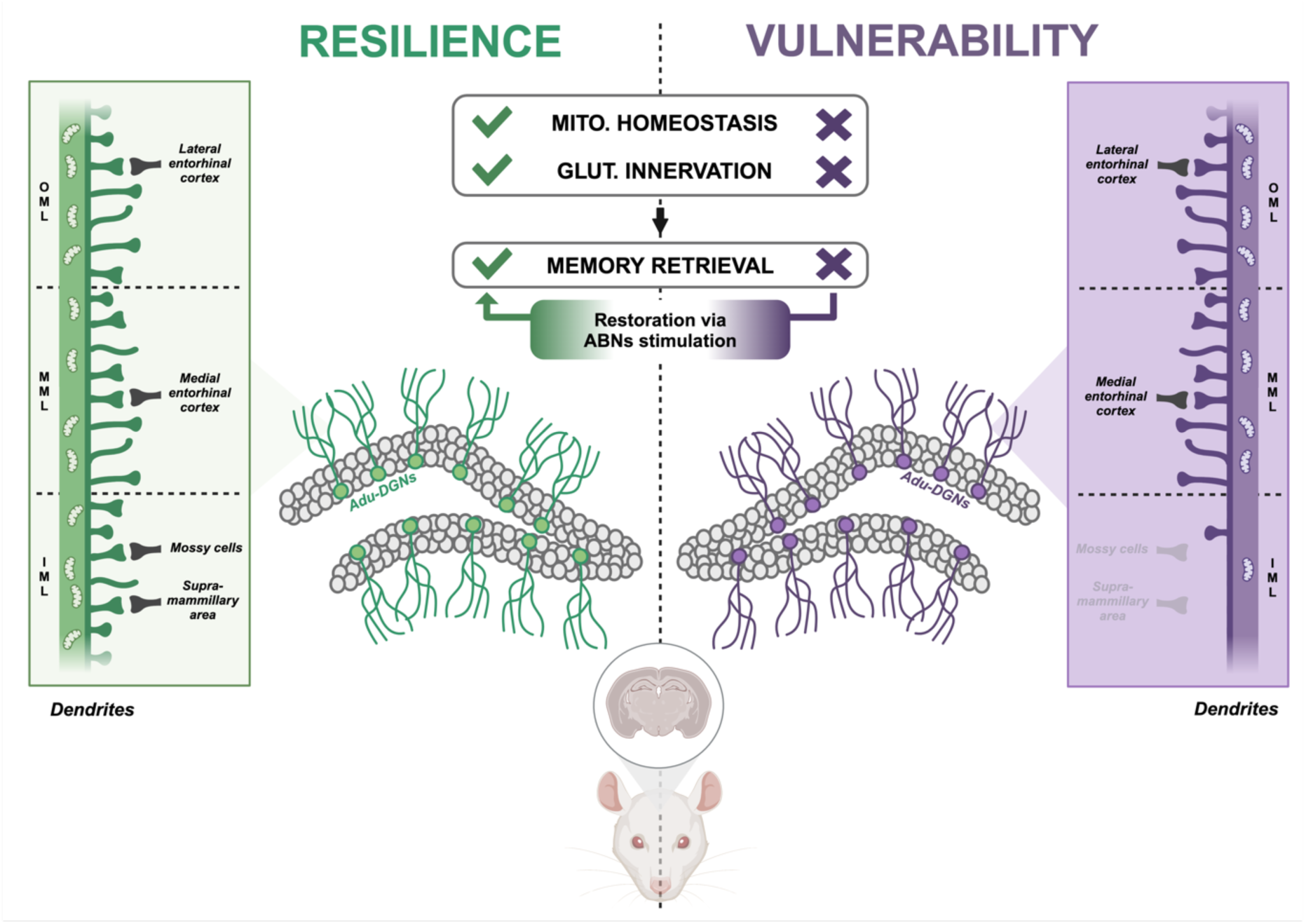

## Main

Cognitive aging has emerged as a global challenge due to the growing elderly population^1^. Cognitive aging is associated with a decline in memory functions, particularly episodic memory which involves recalling personal experiences. Interestingly, this decline varies among individuals, with some remaining resilient with preserved memory functions while others are vulnerable with decreased abilities^2–5^. These inter-individual differences have also been described in rodents^2,3^ especially in tasks measuring spatial memory abilities, making it a good model to study inter-individual differences.

Memory processes depend upon the creation of new neurons in the dentate gyrus of the hippocampus^6,7^. Aging is associated with an exhaustion of the pool of new neurons and their delayed maturation^8–10^. Interestingly, increasing this pool at old age was shown to improve memory abilities^11–14^. However, it is unclear whether neurons born earlier in adult life are also involved in the aging of cognitive functions since most studies focused on neurons produced at old age only. The few data available indicate that these neurons are long-lived as they survive for years and express markers of neuronal activity in response to learning^15,16^.

Here, we tested whether successful aging depends upon the preservation of the health and proper neuronal network integration of mature and long-lived adult-born neurons (ABNs), allowing for accurate reception of incoming information and subsequent recruitment. In contrast, we hypothesized that ABNs in the vulnerable population might experience a progressive disconnection from the network, impeding their recruitment and thereby impairing learning and memory abilities. Thereafter, we investigated whether optogenetics stimulation of ABNs was sufficient to restore memory abilities in aged animals.

### Strategy to follow the aging of ABNs and memory

To analyze the lifelong survival of ABNs born in young adult rats (3-month-old), animals were injected with thymidine analog (Bromodeoxyuridine (BrdU), intraperitoneal injections) that incorporated into dividing cells at the time of injection. To specifically study the morphological features of ABNs, Moloney leukemia virus-based retroviral vectors (M-rv) were injected in the dentate gyrus at the same period. M-rv infect cells dividing at the time of the surgical injections and shortly after^17^. Different M-rv were used to study the morphology (M-rv-GFP^17^), glutamatergic post-synaptic density (M-rv-PSD95-GFP^18^) and mitochondrial network (M-rv-MitoDsRed^19^) of ABNs.

To examine the progressive mechanisms underlying successful aging, we adopted a pseudo-longitudinal strategy in which rats (n=35 per cohort) were submitted to a spatial navigation learning task in the Morris watermaze either at the end of adulthood (8-month-old rats), middle-age (12-month-old rats) or old age (18-month-old rats). This task was chosen as a gold standard method to study cognitive aging as it depends upon adult neurogenesis^20^ and is highly sensitive to aging^3^.

Within each of the three cohorts of rats (adult, middle-aged, old), resilient and vulnerable animals were identified. Subsequently, the morphological analysis was performed in rats with the best memory abilities (n=5, Fig S1), and rats with the worst memory abilities (n=5, Fig S1). Using this strategy, ABNs aged either 5, 9 or 15-month-old in rats aged respectively 8, 12 and 18-month-old at the time of training and sacrifice were analyzed.

### Cell survival and senescence of ABNs do not account for successful cognitive aging

We first examined whether the survival of ABNs tagged in young rats with BrdU was similar or different between the three cohorts of rats of different ages cognitively characterized in the watermaze (Fig1A-B and Fig S1), as it could participate in memory deficits. At all time-points examined, the total number of BrdU labeled cells was similar between resilient and vulnerable animals across all ages (Fig 1C).

**Figure 1:**
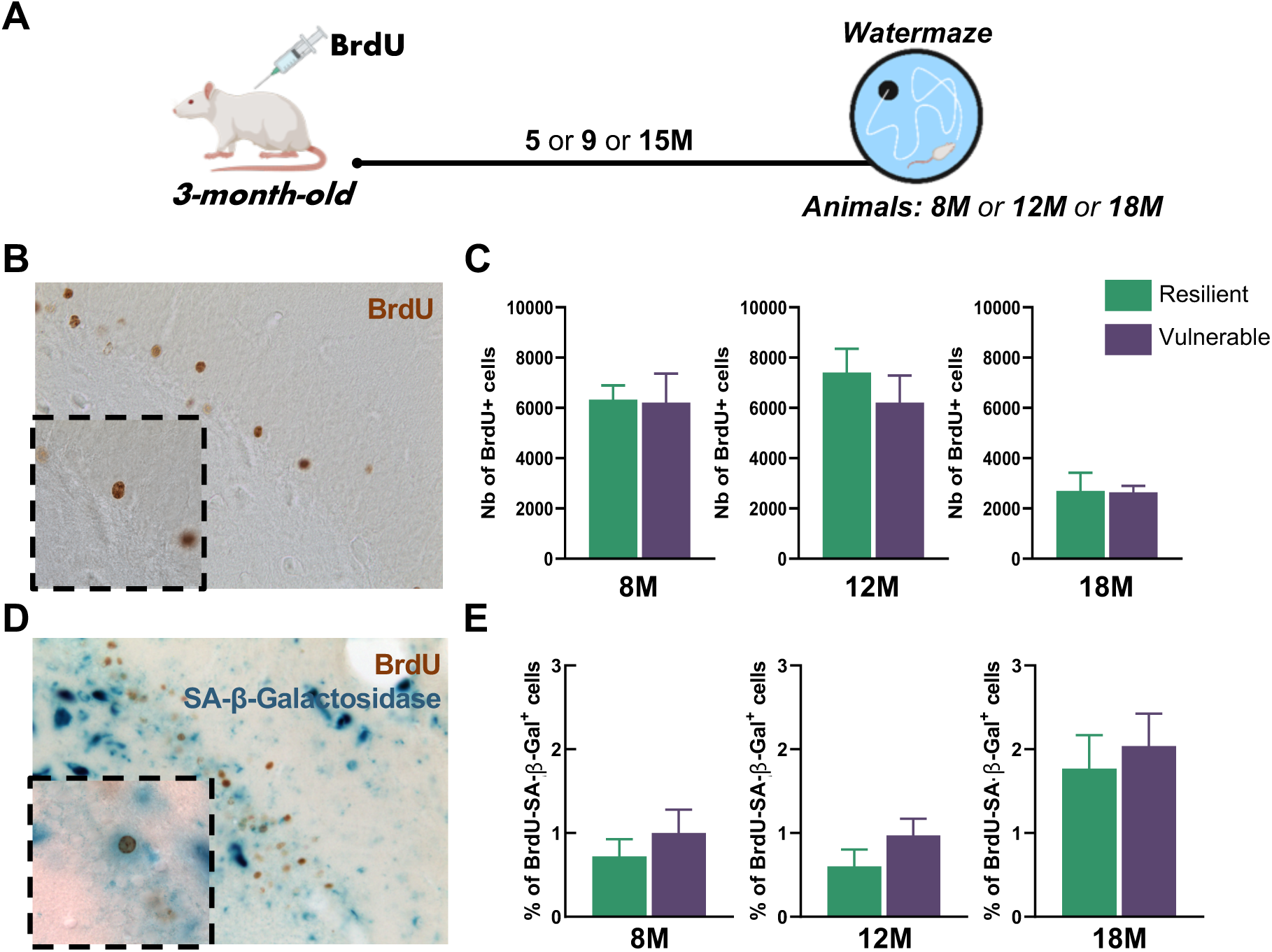
Resilience to cognitive aging is not associated with ABNs death or entry into cellular senescence. (A) Schematic diagram of the experimental design. (B) Picture of a labelled ABNs (Brdu^+^). (C) Resilience to cognitive aging is not linked to different survival of ABNs at 8-month-old (unpaired *t* test: *t_8_* = 0.09, P > 0.05), 12-month-old (unpaired *t* test: *t_8_* = 0.83, P > 0.05) and 18-month-old (unpaired *t* test: *t_8_* = 0.94, P > 0.05). (D) Picture of a senescent ABN (Brdu^+^-SA-β-Gal^+^). (E) Resilience to cognitive aging is not linked to higher levels of senescent ABNs at 8-month-old (unpaired *t* test: *t_8_* = 0.81, P > 0.05); 12-month-old (unpaired *t* test: *t_8_* = 1.29, P > 0.05) or 18-month-old (unpaired *t* test: *t_8_* = 0.48, P > 0.05). Data are presented as mean ± S.E.M. from 5 extreme animals per group. Statistical significance *P ≤ 0.05, **P < 0.01, ***P < 0.001.

Because cellular senescence corresponds to functional cellular arrest and has been described in aged stem cells^21^ and in post-mitotic cells^22^, we used the Senescence-Associated-ß-Galactosidase (SA-ß-Gal) marker to detect entry into senescence of ABNs^23^, as it may account for their reduced recruitment with age. Senescent ABNs were detected and quantified with visualization of cells co-expressing BrdU and SA-ß-Gal (Fig 1D). Resilient and vulnerable animals showed similar levels of senescent ABNs for all ages (Fig 1E). The same profile was observed for the total number of senescent cells (SA-ß-Gal positive) in the granule cell layer of the dentate gyrus (Fig S2). Therefore, while ABNs do not die over the period of 1-2 years, they do not enter into cellular senescence arrest.

### The gross morphological architecture of ABNs is independent of the cognitive status of the rats

One of the hallmarks of neuronal aging is an atrophy of the dendritic network associated with a loss of inputs^24^. To specifically study the morphological features of ABNs, M-rv coupled with GFP (M-rv-GFP^17^) were injected in the dentate gyrus of 3-month-old rats. These M-rv infect ABNs born at the time of the surgery^17^ and allow to label and analyze their somato-dendritic compartments (Fig 2A-2B). Three cohorts of rats of different ages were cognitively characterized (Fig S1), and the total dendritic length of ABNs (Fig 2C), their primary dendrite length (Fig 2D), number of nodes (Fig S3A) and numbers of ends (Fig S3B) were shown to remain similar and independent of the cognitive status of the rats.

**Figure 2:**
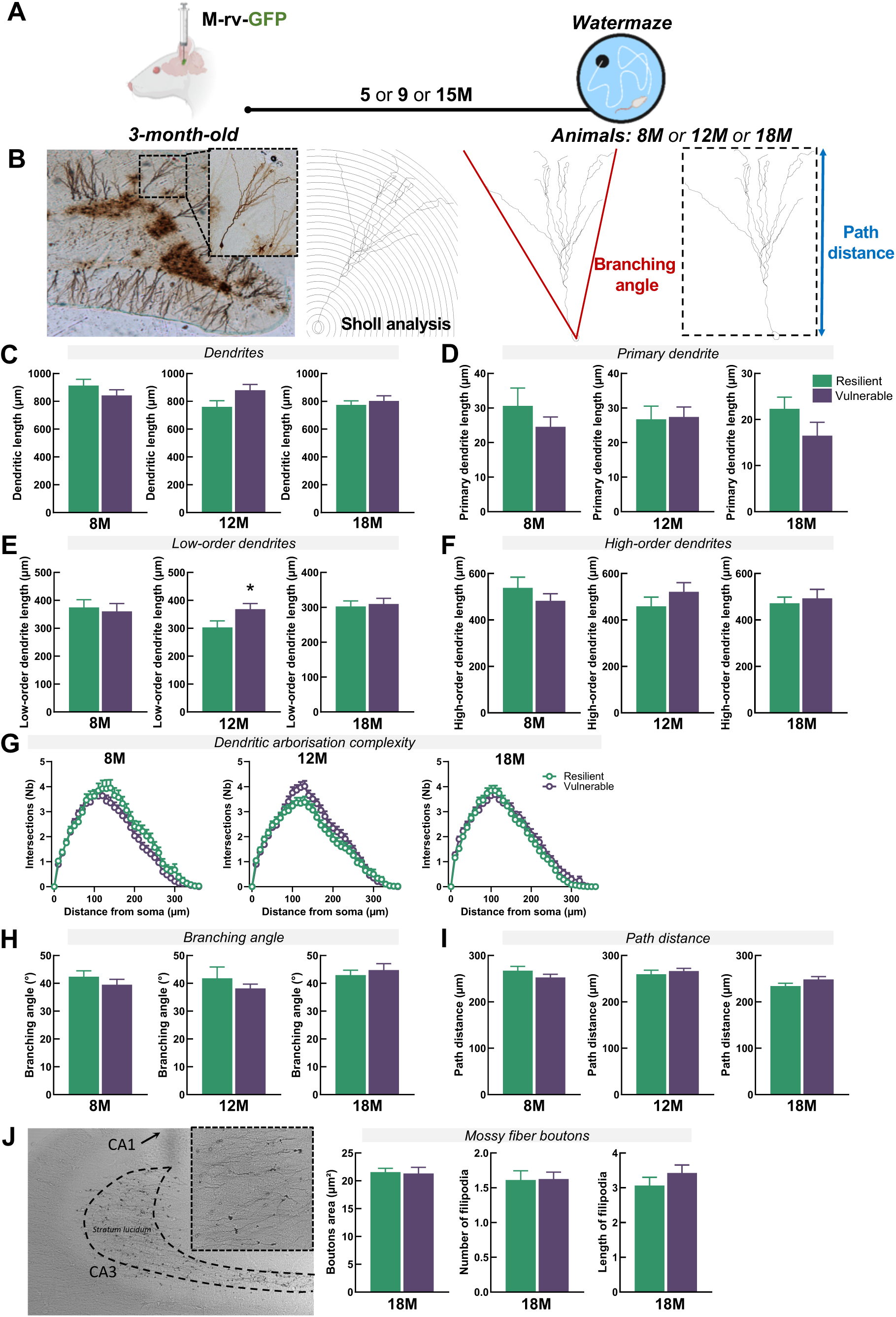
The gross morphological architecture of ABNs is independent of the cognitive status of the rats. (A) Schematic diagram of the experimental design. (B) Picture of a labelled ABN and exemples of Sholl analysis for dendritic complexity, branching angle and path distance calculations. (C) ABNs in the two cognitive populations show similar total dendritic length at 8-month-old (unpaired *t* test: *t_90_* = 1.15, P > 0.05), 12-month-old (unpaired *t* test: *t_102_* = 1.98, P > 0.05) and 18-month-old (unpaired *t* test: *t_111_* = 0.61, P > 0.05). (D) ABNs have similar primary dendritic length in the two cognitive populations at 8-month-old (unpaired *t* test: *t_90_* = 1.11, P > 0.05), 12-month-old (unpaired *t* test: *t_102_* = 0.15, P > 0.05), and 18-month-old (unpaired *t* test: *t_111_* = 1.50, P > 0.05). (E) ABNs have similar low-order dendrites length in the two cognitive populations at 8-month-old (unpaired *t* test: *t_90_* = 0.34, P > 0.05) and 18- month-old (unpaired *t* test: *t_111_* = 0.76, P > 0.05). A small increase in vulnerable animals could be observed at 12-month-old (unpaired *t* test: *t_102_* = 2.12, P ≤ 0.05). (F) ABNs show similar high-order dendrites length between the two cognitive populations at 8-month-old (unpaired *t* test: *t_90_* = 1.11, P > 0.05), 12-month-old (unpaired *t* test: *t_102_* = 0.15, P > 0.05) and 18-month-old (unpaired *t* test: *t_111_* = 1.50, P > 0.05). (G) ABNs of resilient and vulnerable individuals have similar dendritic complexity at 8- month-old (RM-two-way ANOVA, F_36,3240_=1.09, P > 0.05), 12-month-old (RM-two-way ANOVA, F_36,3408_=1.93, P < 0.001) and 18-month-old (RM-two-way ANOVA, F_36,3996_=0.84, P > 0.05). (H) ABNs in the two cognitive populations show similar branching angles at 8-month-old (unpaired *t* test: *t_90_* = 0.99, P > 0.05), 12-month-old (unpaired *t* test: *t_102_* = 0.87, P > 0.05) and 18-month-old (unpaired *t* test: *t_111_* = 0.64, P > 0.05). (I) ABNs show similar path distance between the two cognitive populations at 8- month-old (unpaired *t* test: *t_90_* = 1.30, P > 0.05), 12-month-old (unpaired *t* test: *t_102_* = 0.69, P > 0.05) and 18-month-old (unpaired *t* test: *t_111_* = 1.63, P > 0.05). (J) ABNs show similar MFBs area (unpaired *t* test: *t_28_* = 0.17, P > 0.05), number (unpaired *t* test: *t_28_* = 0.08, P > 0.05) and length (unpaired *t* test: *t_28_* = 1.10, P > 0.05) of filipodia between resilient and vulnerable animals at 18-month-old. Data are presented as mean ± S.E.M. from 5 extreme animal per group (a min of 4 neurons were traced per animal, with 8M-Res = 37 neurons and 8M-Vul = 55 neurons; 12M-Res = 48 neurons and 12M-Vul = 58 neurons; 18M-Res = 63 neurons and 18M-Vul = 50 neurons. Statistical significance *P ≤ 0.05, **P < 0.01, ***P < 0.001.

To better investigate the dendritic complexity, we analyzed the dendrites of ABNs according to their branching order, shown as a good example for sub-dendrite complexity in dentate neurons^25^. However, the ABNs of resilient and vulnerable rats still displayed similar length and number of low and high-order dendrites (Fig 2E-2F and Fig S3C-S3D). Moreover, Sholl analysis revealed that their dendritic arbor had similar complexity (Fig 2G and Fig S3E). In order to further investigate the ABNs integration within the dentate gyrus, the branching angle and path distance of the dendrites were calculated. ABNs in resilient and vulnerable rats displayed similar integration for these parameters as well (Fig 2H-2I). Given this apparent unchanged dendritic morphology, we investigated the morphology of the ABNs outputs at the level of their mossy fiber boutons (MFBs) in the CA3 at the oldest time point (rats aged of 18-month-old). No differences in the MFBs area nor in the number and length of filipodia could be observed, suggesting preserved synaptic outputs in both resilient and vulnerable rats (Fig 2J). Additionally, ABNs in resilient and vulnerable rats showed similar cell body morphologies (Fig S3F). Altogether, no major changes in the gross morphology of ABNs could be evidenced between resilient and vulnerable rats in the course of aging.

### Glutamatergic inputs onto the proximal dendritic segment of ABNs are maintained in resilient rats

Since no major change in the gross morphology of ABNs could be evidenced in the course of aging, we refined our analysis by studying possible connectivity alterations that may lead to altered integration of ABNs within the neuronal network. We focused on the glutamatergic inputs of ABNs, as they represent the main excitatory source essential for neuronal activity and play a key role in spatial learning.

Glutamatergic inputs project to dendritic spines, which are characterized by the presence of the post-synaptic density (PSD), where the PSD95 scaffolding protein is present in high quantity^26^. We used PSD95 as a proxy for glutamatergic inputs onto the spines of ABNs by injecting a M-rv-PSD95-GFP expressing PSD95 coupled with eGFP^18^ (Fig 3A). Since the dentate gyrus receives specific and identified inputs from different glutamatergic regions in each of the sub-layers of the ML [outer molecular layer (OML), middle molecular layer (MML) and inner molecular layer (IML)] (Figure 3B), the quantification of clusters of PSD95 was made for each sub-layer.

**Figure 3:**
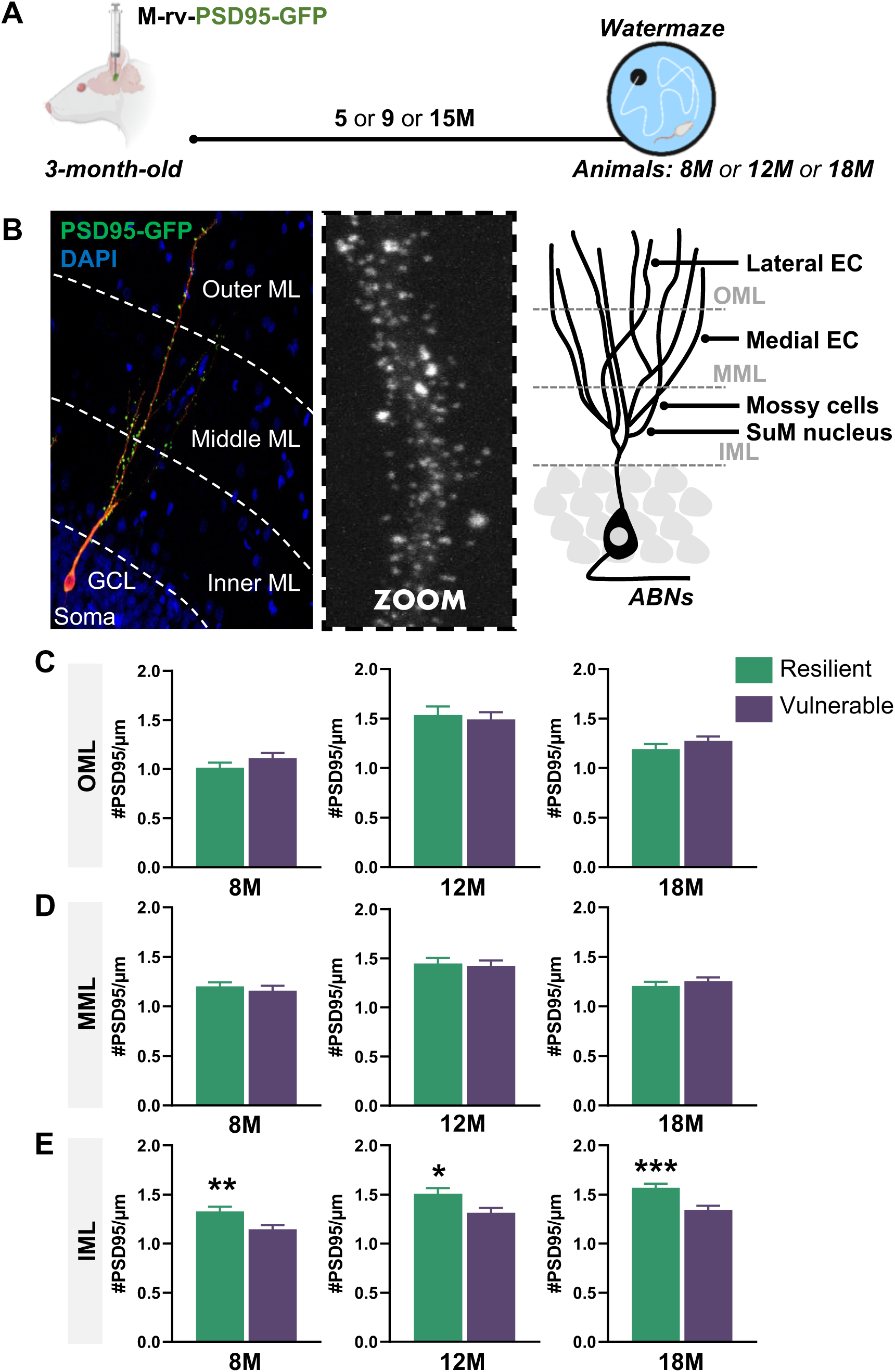
Glutamatergic inputs onto the proximal dendritic segment of ABNs are maintained in resilient rats. (A) Schematic diagram of the experimental design. (B) Picture of a labelled ABN and the subsequent molecular layer (ML) division in three sub-layers; inner (IML), middle (MML) and outer (OML) molecular layer with their different glutamatergic inputs. (C) Resilience to cognitive aging is not linked to the preservation of glutamatergic inputs in the OML at 8-month-old (unpaired *t* test: *t*_104_ = 1.27, P > 0.05), 12-month-old (unpaired *t* test: *t_87_* = 0.38, P > 0.05) or 18-month-old (unpaired *t* test: *t*_154_ = 1.20, P > 0.05). (D) Resilience to cognitive aging is not linked to the preservation of glutamatergic inputs in the MML at 8-month-old (unpaired *t* test: *t_193_* = 0.65, P > 0.05), 12-month-old (unpaired *t* test: *t_199_* = 0.29, P > 0.05) or 18-month-old (unpaired *t* test: *t_210_* = 0.83, P > 0.05). (E) Resilience to cognitive aging is related to the preservation of glutamatergic inputs in the IML at 8-month-old (unpaired *t* test: *t_195_* = 2.73, P < 0.01), 12-month-old (unpaired *t* test: *t_220_* = 2.51, P ≤ 0.05) and 18-month-old (unpaired *t* test: *t_206_* = 3.62, P < 0.001). Data are presented as mean ± S.E.M. from 5 extreme animals per group (a min of 3 neurons were analyzed per animal, with 8M-Res = 27 neurons and 8M-Vul = 27 neurons; 12M-Res = 21 neurons and 12M-Vul = 33 neurons; 18M-Res = 30 neurons and 18M-Vul = 29 neurons. Statistical significance *P ≤ 0.05, **P < 0.01, ***P < 0.001.

For all aged-population cognitively assessed (Fig S1), the ABNs of resilient and vulnerable animals displayed similar post-synaptic densities in the OML (Fig 3C) and MML (Fig 3D). However, ABNs in resilient animals showed a specific preservation of their PSD95 density in the IML compared to ABNs in vulnerable animals for all ages (Fig 3E), suggesting that the specific preservation of the inputs projecting to the proximal dendritic segment of ABNs contributes to resilience in cognitive aging.

### Mitochondrial homeostasis in the proximal dendritic segment of ABNs is maintained in resilient rats

Given that PSD95 is expressed in spines, our previous results suggest that ABNs in the vulnerable population have a reduced number of spines in the IML. Such spine disruption could be linked to metabolic alterations. The mitochondrial network contributes to the formation, maintenance, elimination and functionality of spines^27,28^ and mitochondria depletion was shown to reduce spine density^29^.

According to these observations, we followed the evolution of the mitochondrial network in ABNs with the use of a M-rv-MitoDsRed^19^ (Fig 4A), labelling their mitochondrial network. The quantification of the mitochondrial clusters density along the dendrites was performed in the IML, MML and OML as previously described (Fig 4B). For all cognitively characterized aged-populations (Fig S1), the ABNs of resilient animals exhibited a preserved mitochondrial density in the IML compared to ABNs of vulnerable animals (Fig 4E), corroborating the preservation of the post-synaptic density in the same sub-layer. In the OML and MML, the ABNs in resilient and vulnerable animals showed similar mitochondrial densities at the end of adulthood and middle-age (Fig 4C-4D). However, we observed a spread of the decreased mitochondrial density to these outer layers at old age (Fig 4C-4D). These results indicate that a mitochondrial dysfunction may render vulnerable via the disruption of spines and the subsequent degeneration of inputs projecting to the spines. Therefore, the preservation of mitochondrial homeostasis in ABNs may be essential for successful cognitive aging.

**Figure 4:**
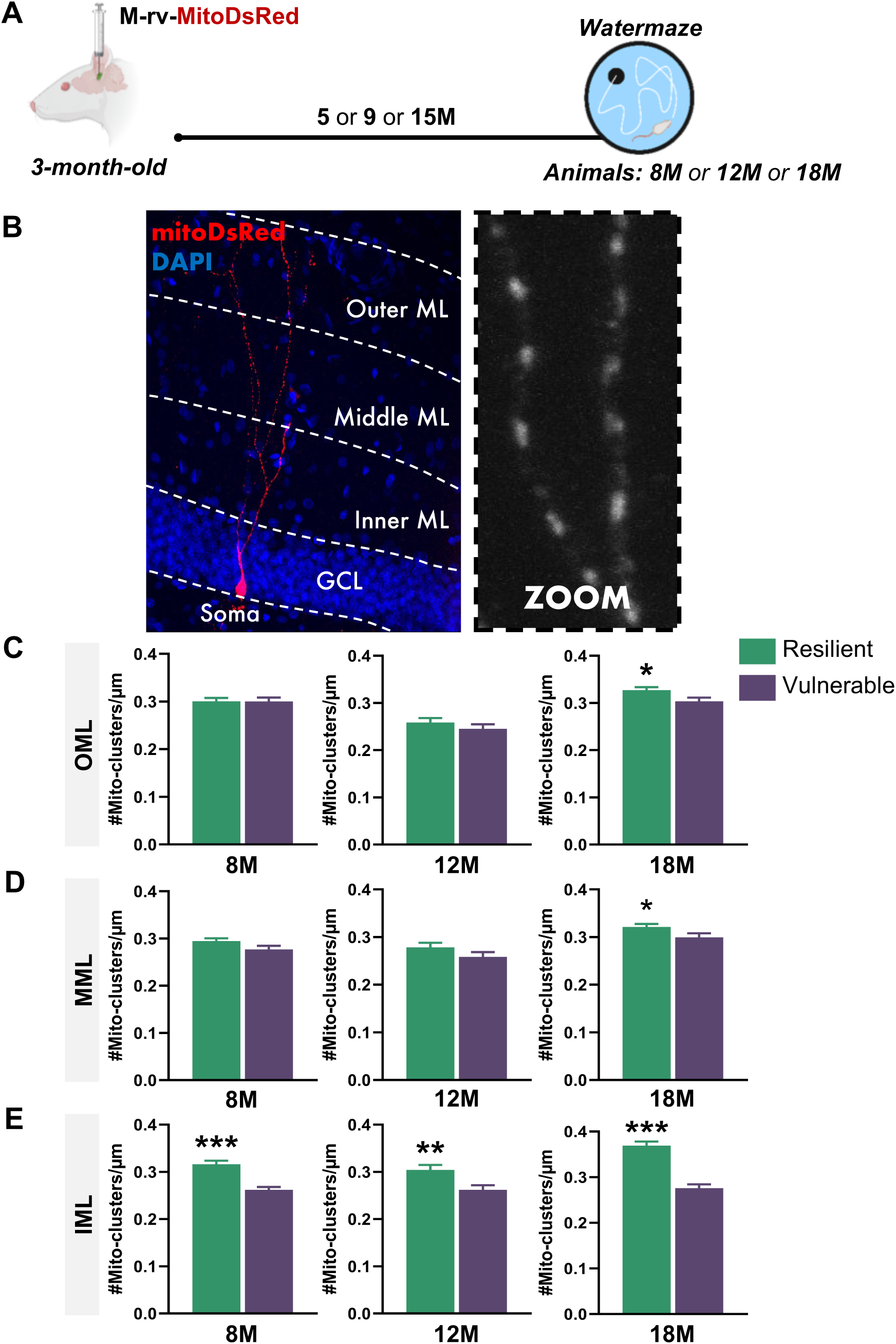
the mitochondrial homeostasis in the proximal segment of dendrites of ABNs is maintained in resilient rats. (A) Schematic diagram of the experimental design. (B) Pictures of labelled ABN and illustatrion of the molecular layer (ML) division in three sub-layers; inner (IML), middle (MML) and outer (OML) molecular layer. (C) Resilience to cognitive aging is linked to the preservation of the mitochondrial density of the OML at 18-month-old (unpaired *t* test: *t_171_* = 2.33, P ≤ 0.05) but not earlier at 8-month-old (unpaired *t* test: *t_154_* = 0.01, P > 0.05) or 12-month-old (unpaired *t* test: *t_134_* = 0.97, P > 0.05). (D) Resilience to cognitive aging is linked to the preservation of the mitochondrial density of the MML at 18-month-old (unpaired *t* test: *t_198_* = 2.01, P ≤ 0.05) but not earlier at 8-month-old (unpaired *t* test: *t_203_* = 1.81, P > 0.05) or 12-month-old (unpaired *t* test: *t_168_* = 1.41, P > 0.05). (E) Resilience to cognitive aging is related to the preservation of the mitochondrial density of the IML at 8-month-old (unpaired *t* test: *t_208_* = 5.53, P < 0.001), 12-month-old (unpaired *t* test: *t_183_* = 2.91, P < 0.01) and 18- month-old (unpaired *t* test: *t_202_* = 7.42, P < 0.001). Data are presented as mean ± S.E.M. from 5 extreme animals per group (a min of 3 neurons were analyzed per animal, with 8M-Res = 28 neurons and 8M-Vul = 25 neurons; 12M-Res = 20 neurons and 12M-Vul = 20 neurons; 18M-Res = 27 neurons and 18M-Vul = 27 neurons). Statistical significance *P ≤ 0.05, **P < 0.01, ***P < 0.001.

### Stimulation of ABNs restores memory retrieval in vulnerable animals

Taken together, the previous data indicate that the observed rarefaction of inputs and disruption of mitochondrial homeostasis could be attributed, at least in part, to a reduction in the functionality of ABNs. To by-pass these alterations in the ABNs network integration, we artificially stimulated ABNs using an optogenetic approach according to a previously used protocol^30^. M-rv-ChannelRhodopsin-GFP was bilaterally injected in the dentate gyrus of 3-month-old rats. At old age (20-month-old), animals were submitted to learning in the watermaze during which ABNs were optogenetically stimulated. We observed similar levels of stimulation of ABNs (aged 17-month-old) between the two aged cognitive populations (Fig S4). Control animals followed the same procedure but without light stimulation. Animals were not over-trained in order to increase the likelihood to enhance the memory trace. Two days after the end of learning, a probe test (during which the platform is removed) was conducted without stimulation in order to test for the strength of memory (Fig 5A). Optogenetic manipulation of ABNs did not improve the learning performances of old animals (Fig S5), as previously shown in younger animals^30^, but improved the memory of the former platform location. We found that the stimulation of ABNs during learning promoted the formation of a precise memory in vulnerable animals, as indicated by the decreased latency to reach the former platform position compared to the non-illuminated vulnerable group, reaching similar latency than for non-stimulated and stimulated resilient animals (Fig 5C). Analysis of the percentage of time spent in the target quadrant (where the escape platform was previously located) revealed that the stimulation of ABNs had further enhanced the memory strength in resilient rats (Fig 5D-5G), as confirmed by the heatmaps analysis of the probe test (Fig 5H). This indicates that optogenetic stimulation of ABNs in vulnerable rats was sufficient to restore memory retrieval to levels observed in non-stimulated resilient animals, but yet not enough to promote the formation of a strong and stable memory trace as it did in stimulated resilient animals. We obtained similar results when the learning and memory test were performed at middle-age (12-month-old) (Fig S6). Altogether, these results indicate that the stimulation of ABNs can restore memory retrieval in vulnerable animals and even further increase the stability and strength of memory in resilient animals.

**Figure 5:**
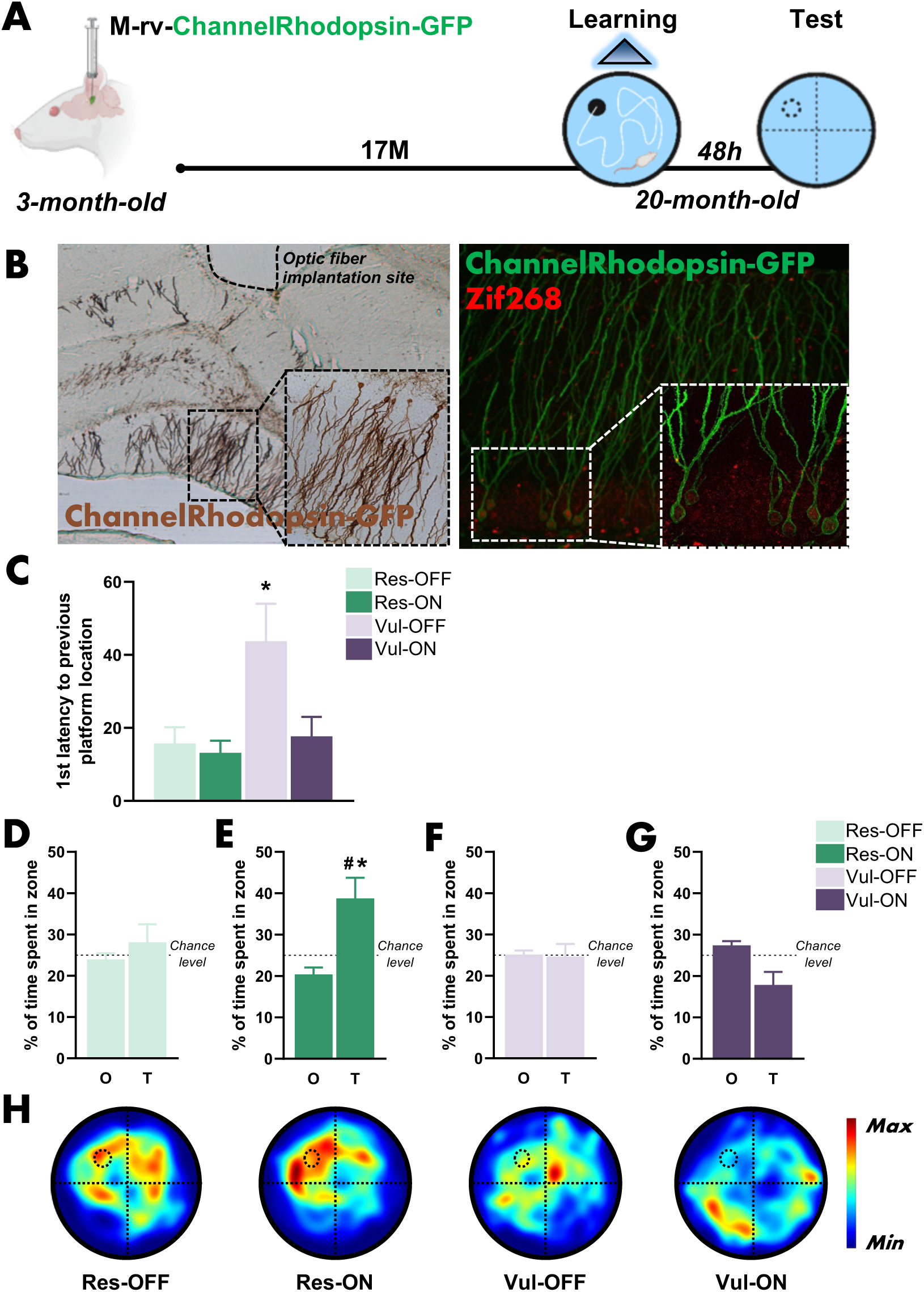
Stimulation of ABNs during learning restores memory retrieval in old vulnerable animals. (A) Schematic diagram of the experimental design. (B) Picture of labelled ABNs (GFP^+)^ and optogenetically stimulated ABNs (GFP^+^-Zif268^+^). (C) Stimulation of ABNs in old stimulated vulnerable (Vul-ON) animals promoted the formation of a precise memory trace similar to resilient animals (Res-OFF and Res-ON) (One-way-ANOVA, F_(3, 13)_ = 5,23, P < 0,05). (D) Old non-stimulated resilient (Res-OFF) animals did not form a stable memory trace (paired *t* test: *t_3_* = 0.72, P > 0.05). (E) Stimulation of ABNs in old stimulated resilient (Res-ON) animals promoted the formation of a strong memory trace (paired *t* test: *t_4_* = 2.77, P ≤ 0.05 and one sample *t* test against ^#^chance level *t*_5_ = 3,23, P ≤ 0.05). (F) Old non-stimulated vulnerable (Vul-OFF) animals did not form a stable memory trace (paired *t* test: *t_3_* = 0.12, P > 0.05). (G) Stimulation of ABNs did not restore the formation of a stable memory trace in Vul-ON animals (paired *t* test: *t_3_* = 2.28, P > 0.05). (H) Heatmaps depicting the animals search location and occupancy during the memory test. Data are presented as mean ± S.E.M. Statistical significance *P ≤ 0.05, **P < 0.01, ***P < 0.001.

## Discussion

Our results highlight the essential role played by long-lived ABNs in successful cognitive aging. By using a novel strategy aiming at tagging ABNs generated during early adulthood, we followed their health and network integration throughout aging. We observed that ABNs in animals with reduced memory abilities progressively lose their glutamatergic innervation in their proximal dendrites, an effect maintained lifelong. This loss of inputs does not appear to be associated with the death or gross morphological deterioration of ABNs. However, we observed a similar reduction of the mitochondrial network density in the proximal dendrites of the ABNs. The reduced mitochondrial density may result from mitochondrial dysfunctions or deposition, which are activity calcium-dependent^31^. Reduced local activity could generate local deposition of mitochondria and hinder their trafficking to the IML. Given that mitochondria represent essential hubs for synaptic functioning and spine maintenance and that mitochondria depletion was shown to reduce spine density^29^, these data suggest a link between the loss of spines (using PSD95 as a proxy) and mitochondria alterations.

We propose that the loss of glutamatergic excitatory innervation, replicated in three independent cohorts of aged rats, could represent the source of the loss of responsiveness of ABNs observed during learning in vulnerable rats^16^. The glutamatergic inputs to the proximal dendrites in the IML come from mossy cells and neurons in the supra-mammillary area (SuM) of the hypothalamus. Very little is known about these two types of projections to the dentate gyrus in the context of inter-individual differences in cognitive aging. However, studies have shown that mossy cells^32–34^ and the SuM^35,36^ participate in spatial memory retrieval, and the disruption of their projection to the dentate gyrus may result in the cognitive deficits we observed. Therefore, we have identified for the first time that mossy cells and the SuM are two new potential players in resilience to cognitive aging appearing as early as middle age.

To date, only the MML and OML connectivity of the dentate gyrus had been addressed in the context of inter-individual differences in cognitive aging. Using electronic microcopy, the number of axo-spinous synapses was shown to be reduced in the MML of 24-28-month-old rats exhibiting memory deficits^37^. In the same line of idea, the synaptophysin levels, a marker for pre-synaptic vesicles, indicated impaired connectivity in the MML and OML of vulnerable rats^38^. These results suggested that resilience to cognitive aging is linked to the preservation of inputs from the entorhinal cortex. While we did not observe differences in the post-synaptic density of MML and OML, we observed the emergence of alterations in the ABNs mitochondrial network in these outer layers in 18-month-old vulnerable rats (Fig 4C-4D). As previously proposed, mitochondrial disruptions might lead to spine disruption and to the loss of inputs on the former spine. Given that the previous studies addressed the dentate connectivity in older rats aged 24-28-month-old^37,38^, it likely allowed them to identify alterations that we might have detected with our PSD95 tool at timepoints beyond 18-month-old in the MML and OML. Nevertheless, we cannot exclude that the tool used in our present study is more coarse-grained than electronic microscopy, suggesting that we may have missed some changes at the level of inputs from the entorhinal cortex. In addition, we cannot exclude the involvement of other inputs. For example, cholinergic inputs from the medial septum projects on dentate neurons and make asymmetric and presumably excitatory contacts on dendritic spines, chiefly in the IML^39^, and are known to degenerate in vulnerable rats^40^.

We propose that resilience to cognitive aging in our behavioral paradigm specifically depends on ABNs, as opposed to other dentate neuron generated during development. Indeed, dentate neurons born during development are not needed for spatial learning^30^ or memory retrieval and reconsolidation^41,42^ and are not recruited by spatial learning in both young adults^43^ and old individuals^16^. Consequently, the observed loss of glutamatergic inputs from mossy cells and the SuM in the proximal dendrites should be specific to ABNs. To test this hypothesis, we quantified the total glutamatergic innervation in the IML, MML and OML in our oldest cohort of animals (18 months old) using vGLUT2 labelling, a marker for presynaptic glutamatergic terminals^44^. We also quantified the total number of mossy cells in the dentate hilus. Since developmentally-born neurons are significantly more abundant in the dentate gyrus compared to ABNs, it is more likely that these quantifications correspond to developmentally-born neurons^45,46^. We observed no differences in the dentate glutamatergic innervation across each sub-layer of the ML using vGLUT2 labelling (Fig S7), nor in the total number of mossy cells (Fig S8), regardless of the cognitive status at old age. Based on these observations, we propose that in our behavioral paradigm resilience to cognitive aging primarily relies on ABNs and their ability to maintain proper and functional integration into the neuronal network and circuitry during aging.

The differences observed in glutamatergic innervation and mitochondrial density of ABNs might originate from the dysregulation of the hypothalamic–pituitary–adrenal (HPA) axis in vulnerable rats. Indeed, animals displaying high-reactivity to stress and a hyper-active HPA axis were shown to be prone to develop age-related memory impairments^12,47,48^. Besides, over-exposition to glucocorticoid stress hormones were also shown to be linked to reduced levels of neurogenesis at old age. Interestingly, suppressing glucocorticoid exposition from middle-age has been shown to successfully prevent the emergence of age related memory disorder^12^. Given that glucocorticoid exposition has also been shown to disturb synaptic density and mitochondrial homeostasis and trafficking^49,50^, it could be suggested that an early hyper-activity of the HPA axis in vulnerable individuals (and therefore over-exposition to glucocorticoids) results in our observations. Along the brain–body axis, an early alteration in the aging of the systemic milieu could also be at play^51^, as blood circulating factors have been shown to affect adult neurogenesis^52–54^ and so did secreted factors such as irisin^55^ or osteocalcin^56^.

Interestingly, despite alterations observed as early as 8-month-old in animals, we successfully restored memory retrieval abilities in middle-age (12-month-old) and old (18-month-old) vulnerable rats using optogenetic stimulation of long-lived ABNs. This suggests that ABNs remain functionally viable in vulnerable rats and can transmit information downstream in the network for memory processing, as evidenced by their preserved output connectivity, and that the primary obstacle to their recruitment appears to be the rarefaction of their inputs. While we succeeded in restoring the memory retrieval abilities of stimulated vulnerable animals to levels comparable to non-stimulated resilient animals, it is noteworthy that this intervention did not establish a strong and lasting memory trace as observed in stimulated resilient animals. This indicates that optimal memory performance can be further achieved by stimulating neurons that are well-integrated within the neuronal network. Therefore, future therapeutic approaches aimed at restoring memory functions should target long-lived ABNs, as they clearly still participate in memory processes, a fact often overlooked in the literature. However, it is crucial to address the alteration of inputs detected as early as middle-age and to aim at preventing it to achieve optimal therapeutic results in old age.

In conclusion, our results indicate that brain resilience relies on the preservation of the integration of ABNs into the network and circuitry. We highlighted the essential role of long-lived ABNs in resilience against cognitive decline and revealed their potential as a valuable tool for rejuvenating memory abilities at old age.

### Material & methods

#### 1. Rats

A total number of 281 male Sprague-Dawley rats (OFA, Janvier, France) were used for these experiments. Rats aged 3-month-old at the time of delivery were grouped housed in standard cages under a 12/12h light/dark cycle with ad libitum access to food and water. Temperature (22°C) and humidity (60%) were kept constant. Rats were individually housed before the beginning of training at either the end of adulthood (8-month-old), middle-age (12-month-old) or old age (18 or 20-month-old). Experimental procedures have been carried out following the European directive of the parliament and the conceal of September 22, 2010 (2010/63/UE). Animal studies were approved by the ethical committee of the University of Bordeaux (ID #10864, #11155, #21501, #5012006A).

#### 2. Thymidine analog injections

BrdU (5-bromo-2′-deoxyuridine) and CldU (5-chloro-2’-deoxyuridine) were dissolved in a Phosphate Buffer (pH 8.4) and NaCl, respectively. CldU injections were equimolar doses of BrdU (50mg/kg). Rats received four injections intra-peritoneally during four consecutive days when 3-month-old.

#### 3. Retroviruses

The M-rv-GFP has been described previously^17^ and is a gift from Prof. F.H. Gage (Salk Institute, La Jolla, CA). The M-rv-PSD95-GFP has been described previously^18^ and is a gift from Dr C. Lois (Caltech, Pasadena, CA). The M-rv-MitoDsRed has been described previously^19^ and is a gift from Dr D.C. Lie (Friedrich-Alexander Universität Erlangen-Nürnberg, Erlangen, Germany). The M-rv-Channelrhodopsin-GFP was kindly provided by the Dr S. Ge (Stony Brook University, Ney York City, NY). High retroviruses titles (between 5 × 10^8^ and 5 × 10^9^) were prepared with a human 293-derived retroviral packaging cell line (293GPG) kindly provided by Dr D.C. Lie. Virus-containing supernatant was harvested three days after transfection with Lipofectamine 2000 (Thermofisher). This supernatant was then cleared from cell debris by centrifugation at 3500 rpm for 15 min and filtered through a 0.45 μm filter (Millipore). Viruses were concentrated by two rounds of centrifugation (19 500 rpm 2 h) and resuspended in PBS.

#### 4. Surgery

Rats were anaesthetized with 3% isoflurane and placed in the stereotaxic frame, where they were maintained asleep with 2% isoflurane for the duration of the surgery. Analgesia was provided by an injection of Metacam (1ml/kg, sub-cutaneously, Boehringer Ingelheim) and lidocaine (0.1 ml, sub-cutaneous, Vetoquinol) at the incision site.

##### Retroviral injections

A retroviral mix of half M-rv-GFP and half M-rv-MitoDsRed was prepared before injection. The retroviral injections (2 μL per injection site at 0.3 μL/min) were stereotaxically made into the dentate gyrus of 3-month-old adult rats with a microcapillary pipette connected to a Hamilton syringe placed into a micro injector (KDS legato 130) directly attached to the stereotaxic frame. Cannulas were maintained in position for 3 minutes after the end of the injections to let the suspension infuse. The Moloney leukemia virus-based retroviral vector M-rv-PSD95-GFP and mixed M-rv-GFP/M-rv-MitosDsRed were injected to the right and left dorsal dentate gyrus respectively. For the optogenetic experimental cohorts, the M-rv-Channelrhodopsin-GFP was injected bilaterally to the dorsal area of the dentate gyrus. Injections were administered using the following coordinates: [AP]:-3,5mm; [L]:- /+1,7mm; [P] from skull:-4,3mm.

##### Optic fiber implantations

Home-made optic fibers were made following a protocol previously described^57^. The implantation surgical procedure was carried out one month before the start of behavior, following the same protocol as that described for the retroviral injection. The skull was scratched and three anchor screws were fixed (one in the anterior area and two in the posterior area). The two optic fibers [length: 4.5 mm, diameter: 200 μm, numeric aperture (NA: 0.37)] were lowered with a speed of ∼2 mm/min upwards to the M-rv injection site with the following stereotaxic coordinates: [AP]:-4,1mm; [L]:-/+1,8mm; [P] from *dura-mater*:-3mm. Fibers were stabilized using dental cement. When the cement was dry, scalps were sutured and disinfected with local antiseptic treatment (Betadine). All rats were followed up until the start of the behavior. During handling, fur was checked and the scar disinfected systematically. Rats were handled every day to habituate them before the behavioral procedure.

#### 5. Watermaze procedures

The apparatus consisted of a circular plastic swimming pool (180 cm diameter, 60 cm height) filled with water (20 ± 1 °C) rendered opaque by the addition of a white cosmetic adjuvant. Two days before training, the animals were habituated to the pool for 1 min. For old animals, we followed previous protocol published for aging experiments in the laboratory^58^. The capacity to swim and to see correctly were assessed during a training phase where animals had to locate a visible platform within the swimming pool, all rats with deficits were removed from the experiments (i.e. rats that did not find the platform in less than 20s after 5 day of platform training). During training, animals were required to locate a submerged platform hidden 1.5 cm under the surface of the water in a fixed location, using the spatial cues available in the room. Rats were trained for four trials per day (90 s with an inter-trial interval of 30 s and released from three different starting points that varied randomly each day). If an animal failed to locate the platform, it was placed on the platform at the end of the trial for the duration of the inter-trial interval. The distance crossed to reach the platform was recorded using a video camera that was secured to the ceiling of the room and connected to a computerized tracking system (Videotrack, Viewpoint). Daily results were analyzed in order to rank animals according to their behavioral score calculated over the last 3 days of training. Training stopped before the resilient animals reached plateau phase of learning.

##### Watermaze for optogenetic manipulation of ABNs

The watermaze was equipped with a blue laser source (OptoDuet 480 nm, 200mW, Ikecool) connected to a rotatory joint to avoid tangling of the patch cords connected to the intra-dentate gyrus optic fibers. Light intensity at the optic fiber cable tip was controlled every day before the beginning of the session (8-10 mV). Illumination was applied throughout the session at 20 Hz pulses of 15 ms. The time to reach the platform was recorded using a video camera connected to a computerized tracking system (Polyfiles, Imetronic). Light stimulation was provided during the 4 trials each day for the entire learning phase. When resilient animals reached the plateau phase, training was stopped and 48h later animals were tested during a probe test for 60 s where no animals were under no-light stimulation. During the probe test, the hidden platform was removed and the time spent in each quadrant was quantified. Control animals consisted of animals that underwent the same procedure but without light stimulation.

###### Resilient and vulnerable extremes populations

Following the end of learning, animals were cognitively attributed according to the mean performances of their three last days of training. From the whole population submitted to the learning task, the five animals with the higher score (i.e. lowest mean distance to reach the platform) were attributed as extreme resilient and the five animals with the lowest score (higher mean distance to reach the platform) were attributed as extreme vulnerable. Only the extremes were kept for analysis as considered the most representative of their cognitive abilities, as already published in the laboratory^58^.

### 6. Immunohistochemistry and analysis

For animals injected with M-rv-GFP, M-rv-MitoDsRed, M-rv-PSD95-GFP and M-rv-Channelrhodospin-GFP, animals were perfused transcardially with a phosphate-buffered solution of 4% paraformaldehyde 90 min after the last training session. After 24h fixation, optogenetic brains were cut using a vibratome. For PSD95-GFP and mitoDsRed, after 24h of fixation, brains were transferred in a PBS-sucrose (30% solution for 24h and then cut using a vibratome.

#### GFP expressing cells

Free-floating 50-μm-thick sections were processed according to a standard immunohistochemical procedure on alternate one-in-ten sections to visualize the retrovirus-expressing GFP with an anti-GFP primary antibody (Rabbit, 1:12000; Millipore, Cat#AB3080P). Bound antibodies were visualized using the biotin-streptavidin technique (ABC kit, Vector Laboratories Inc., Cat#PK-4000) and 3,3′-diaminobenzidine as a chromogen with a biotinylated goat anti-rabbit antibody (1:200, Vector Laboratories Inc., Cat#BA-1000). Primary antibody was incubated at 4°C for 72h, and secondary antibodies was incubated at room temperature (RT) for 2h. The morphometric analysis of virus-labelled neurons was performed with a ×100 objective. Measurements of dendritic parameters as well as Sholl analysis were performed with the Neurolucida (software Microbrightfield, Colechester, VT, USA). The analysis of the dendritic arborization (low and high order repartition, branching angle and path distance calculations) was performed as previously described^25^ blind by the person who carried it out. Only animals with at least 4 neurons who could be reconstructed and neurons with a minimum of 4 ramifications point were kept for analysis. The analysis was performed blind by the person who carried it out.

#### PSD95-GFP and mitoDsRed expressing cells

Free-floating 50-μm-thick sections were processed according to standard immunohistofluorescence procedure on alternate one-in-ten sections to visualize the retrovirus-expressing eGFP and mitoDsRed with an anti-GFP primary antibody (Rabbit, 1:500; Millipore, Cat#AB3080P) and anti-DsRed primary antibody (Rabbit, 1:1000, Clontech Takara, Cat#632496). Bound antibodies were visualized with with Cy3 goat anti-rabbit (Jackson Immuno Research, 1:1000, Cat#111-167-003) secondary antibody. Primary antibodies were incubated simultaneously at 4°C for 72h, and secondary antibodies were incubated simultaneously at room temperature (RT) for 2h. Fluorescence was studied using a SPE confocal system with a plane apochromatic X 63 oil lens (numerical aperture 1.4; Leica) and a digital zoom of 2,5. For both approaches, MosaicJ plugin from ImageJ was used to reconstruct the neuron as a whole. Mosaic reconstruction allowed to determine the length of the molecular layer and to delimitate the inner, middle and outer sub-layers of the molecular layer. For each layer, dendritic segments of minimum 15 µm and maximum 50 µm were analyzed (minimum of 3 neurons per animals, 5 animals per group). The number of clusters of PSD95-GFP and mitoDsRed was determined either manually or semi-automatically using an intensity threshold in Image J. The density of clusters was obtained by dividing the number of clusters by the length of the corresponding dendritic segment. The analysis was performed blind by the person who carried it out.

##### BrdU and Senescence-Associated-ß-Galactosidase (SA-ß-Gal) positive cells

BrdU^+^ cells throughout the entire granular layer of the dentate gyrus were revealed using the biotin-streptavidin technique with a horse anti-mouse antibody (1:200, Vector Labs, Peterborough, UK. #BA-2001). Primary antibody was incubated at 4°C for 72h, and secondary antibodies was incubated at room temperature (RT) for 2h. SA-ß-Gal^+^ cells were revealed using the Senescence β-Galactosidase Staining Kit (Cell Signaling, #9860) according to the manufacturer’s instructions. The number of BrdU^+^ cells and SA-ß-Gal^+^ cells in the dentate gyrus was estimated with systematic random sampling of every tenth section along the septo-temporal axis of the hippocampal formation using a modified version of the optical fractionator method. Indeed, all of the BrdU-SA-ß-Gal^+^ cells were counted on each section and the resulting numbers were tallied and multiplied by the inverse of the section sampling fraction (1/ssf = 10). The percentage of colocalization BrdU-SA-ß-Gal^+^ cells was calculated as follows: (Nb of BrdU^+^-SA-ß-Gal^+^) / (Nb of BrdU^+^-SA-ß-Gal^+^ + Nb of BrdU^+^-SA-ß-Gal^-^) x 100. The analysis was performed blind by the person who carried it out.

### 7. Statistical analysis

The data (mean ± SEM) were analyzed using the Student *t*-test (two-tailed, paired or unpaired when appropriate) or one way and repeated-measures two-way ANOVA, then followed by Tukey’s comparison test when necessary. Data were tested for normality. All analyses were carried out using the software GraphPad Prisms 8.

## Supplementary Material & methods

### Channelrhodospine-GFP expressing cells

Free-floating 50-μm-thick sections were processed according to a standard immunohistochemical procedure on alternate one-in-ten sections to visualize the retrovirus-expressing GFP with an anti-GFP primary antibody (Rabbit, 1:2000; Millipore, Cat#AB3080P). Primary antibody was incubated at 4°C for 72h, and secondary antibodies was incubated at room temperature (RT) for 2h. Bound antibodies were visualized using the biotin-streptavidin technique (ABC kit, Vector Laboratories Inc., Cat#PK-4000) and 3,3′-diaminobenzidine as a chromogen with a biotinylated goat anti-rabbit antibody (1:200, Vector Laboratories Inc., Cat#BA-1000). Channelrhodopsin-GFP^+^ cells were counted under a ×100 microscope objective throughout the entire septotemporal axis of the granule layer of the dentate gyrus. The total number of cells was estimated using the optical fractionator method, and the resulting numbers were tallied and multiplied by the inverse of the sections sampling fraction (1/ssf10). The analysis was performed blind by the person who carried it out.

### Channelrhodospine-GFP and Zif268 colocalisation

The activation of Channelrhodopsine-GFP^+^ cells was studied using immunohistofluorescence procedure on alternate one-in-ten sections. The retrovirus-expressing GFP was visualized with an anti-GFP primary antibody (Chicken-1:1000, Abcam, Cat#ab13970) and cellular activity with an anti-Zif268 antibody (Rabbit-1:100, Santa Cruz Biotechnology, Cat#sc-189). Bound antibodies were visualized with Alexa488 goat anti-chicken (Thermo Fisher, 1:1000, Cat#A-11039) and Cy3 goat anti-rabbit (Jackson Immuno Research, 1:1000, Cat#111-167-003) secondary antibodies. Primary antibodies were incubated simultaneously at 4 °C for 72h, and secondary antibodies were incubated simultaneously at RT for 2h. Fluorescence was studied using a SPE confocal system with a plane apochromatic X 63 oil lens (numerical aperture 1.4; Leica) and a digital zoom of 2,5. The percentage of GFP cells expressing IEG (all along the temporal–septal axis) was calculated as follows: (Nb of GFP^+^-IEG^+^ cells)/(Nb of GFP^+^-IEG^−^ cells + Nb of GFP^+^-IEG^+^ cells) × 100. In middle-age experiments, every GFP^+^ were analyzed in the left and right dentate gyrus. In old age experiments, every GFP^+^ cells were analyzed in the left dentate gyrus. The analysis was performed blind by the person who carried it out.

### vGLUT2 labelling

Free-floating 50-μm-thick sections were processed according to standard immunohistofluorescence procedure on alternate one-in-ten sections to visualize vGLUT2 in pre-synaptic terminals with an anti-vGLUT2 primary antibody (Rabbit, 1:250; Synaptic Systems, Cat#135403). Bound antibodies were visualized with Cy3 goat anti-rabbit (Jackson Immuno Research, 1:1000, Cat#111-167-003) secondary antibody. Primary antibody was incubated at 4°C for 72h, and secondary antibody was incubated at room temperature (RT) for 2h30. Fluorescence was studied using a SPE confocal system with a plane apochromatic X 20 oil lens (numerical aperture 1.4; Leica) and a digital zoom of 1. Acquisitions were performed in the dorsal dentate gyrus followed by the reconstruction of the acquired dentate gyrus using mosaics, resulting in 3 mosaic-reconstructed dentate gyrus analyzed per animal. An Image J macro was used to delimitate the inner, middle and outer sub-layers of the molecular layer on the mosaic and to calculate the mean intensity of vGLUT2 fluorescent labelling. The mean intensity for vGLUT2 labelling was obtained by dividing the mean intensity of each area analyzed (mean intensity / µm2). The analysis was performed blind by the person who carried it out.

### Mossy cells GluR2/3 labelling

Free-floating 50-μm-thick sections were processed according to a standard immunohistochemical procedure on alternate one-in-ten sections to visualize GluR2/3 labelling expressed in mossy cells with an anti-GluR2/3 primary antibody (Rabbit, 1:50; Millipore, Cat#AB1506). Primary antibody was incubated at 4°C for 72h, and secondary antibodies was incubated at room temperature (RT) for 2h. Bound antibodies were visualized using the biotin-streptavidin technique (ABC kit, Vector Laboratories Inc., Cat#PK-4000) and 3,3′-diaminobenzidine as a chromogen with a biotinylated goat anti-rabbit antibody (1:200, Vector Laboratories Inc., Cat#BA-1000). The total number of mossy cells were counted under a X 40 microscope objective all along the temporal–septal axis of the left and right dentate gyrus. The total number of cells was estimated using the optical fractionator method, and the resulting numbers were tallied and multiplied by the inverse of the sections sampling fraction (1/ssf10). The analysis was performed blind by the person who carried it out.

## General

The authors gratefully acknowledge C. Dupuy for animal care and Léa Archi, Murielle Fèvre, Anaïs Pochat and Loreen Rupprecht for their participation on the work. The authors acknowledge Dr. Fred H. Gage (Salk Institute – La Jolla – CA) and Dr. Shaoyu Ge (Stony Brook University – New York - NY) for the gift of viral-vectors (GFP and Channelrhodopsin-GFP). The confocal analysis was done in the Bordeaux Imaging Center (BIC), a service unit of the CNRS-INSERM and Bordeaux University, member of the National Infrastructure France BioImaging. Figures were created using BioRender with a full license authorization to publish.

## Authors contribution

CL and DCL provided the retroviral constructs. FF produced the retroviruses. NB and VC designed, performed and analyzed the experiments. EC analyzed the mossy fiber boutons. NB and NM performed the optogenetic experiments. EP and MK advised as experts. NB and DNA wrote the manuscript. NB, VC, EP, MK, DCL and DNA revised and approved the manuscript. DNA conceived and designed experiments and provided funding.

## References

1. World Population Prospects 2022: Summary of Results. (United Nations, New York, 2022).

2. Rapp, P. R. & Amaral, D. G. Individual differences in the cognitive and neurobiological consequences of normal aging. Trends Neurosci. 15, 340–345 (1992).

3. Gallagher, M., Burwell, R. & Burchinal, M. Severity of spatial learning impairment in aging: development of a learning index for performance in the Morris water maze. Behav. Neurosci. 107, 618– 626 (1993).

4. Gallagher, M. & Rapp, P. R. The use of animal models to study the effects of aging on cognition. Annu. Rev. Psychol. 48, 339–370 (1997).

5. Nyberg, L., Lövdén, M., Riklund, K., Lindenberger, U. & Bäckman, L. Memory aging and brain maintenance. Trends Cogn. Sci. 16, 292–305 (2012).

6. Anacker, C. & Hen, R. Adult hippocampal neurogenesis and cognitive flexibility - linking memory and mood. Nat. Rev. Neurosci. 18, 335–346 (2017).

7. Abrous, D. N., Koehl, M. & Lemoine, M. A Baldwin interpretation of adult hippocampal neurogenesis: from functional relevance to physiopathology. Mol. Psychiatry 27, 383–402 (2022).

8. Drapeau, E. & Nora Abrous, D. Stem Cell Review Series: Role of neurogenesis in age-related memory disorders. Aging Cell 7, 569–589 (2008).

9. McAvoy, K. M. & Sahay, A. Targeting Adult Neurogenesis to Optimize Hippocampal Circuits in Aging. Neurother. J. Am. Soc. Exp. Neurother. 14, 630–645 (2017).

10. Trinchero, M. F. et al. High Plasticity of New Granule Cells in the Aging Hippocampus. Cell Rep. 21, 1129–1139 (2017).

11. Kempermann, G., Kuhn, H. G. & Gage, F. H. Experience-induced neurogenesis in the senescent dentate gyrus. J. Neurosci. Off. J. Soc. Neurosci. 18, 3206–3212 (1998).

12. Montaron, M. F. et al. Lifelong corticosterone level determines age-related decline in neurogenesis and memory. Neurobiol. Aging 27, 645–654 (2006).

13. McAvoy, K. M. et al. Modulating Neuronal Competition Dynamics in the Dentate Gyrus to Rejuvenate Aging Memory Circuits. Neuron 91, 1356–1373 (2016).

14. Berdugo-Vega, G. et al. Increasing neurogenesis refines hippocampal activity rejuvenating navigational learning strategies and contextual memory throughout life. Nat. Commun. 11, 135 (2020).

15. Kempermann, G., Gast, D., Kronenberg, G., Yamaguchi, M. & Gage, F. H. Early determination and long-term persistence of adult-generated new neurons in the hippocampus of mice. Dev. Camb. Engl. 130, 391–399 (2003).

16. Montaron, M., Charrier, V., Blin, N., Garcia, P. & Abrous, D. N. Responsiveness of dentate neurons generated throughout adult life is associated with resilience to cognitive aging. Aging Cell 19, (2020).

17. Zhao, C., Teng, E. M., Summers, R. G., Ming, G. & Gage, F. H. Distinct Morphological Stages of Dentate Granule Neuron Maturation in the Adult Mouse Hippocampus. J. Neurosci. 26, 3–11 (2006).

18. Kelsch, W., Lin, C.-W. & Lois, C. Sequential development of synapses in dendritic domains during adult neurogenesis. Proc. Natl. Acad. Sci. U. S. A. 105, 16803–16808 (2008).

19. Steib, K., Schäffner, I., Jagasia, R., Ebert, B. & Lie, D. C. Mitochondria modify exercise-induced development of stem cell-derived neurons in the adult brain. J. Neurosci. Off. J. Soc. Neurosci. 34, 6624–6633 (2014).

20. Dupret, D. et al. Spatial Relational Memory Requires Hippocampal Adult Neurogenesis. PLoS ONE 3, e1959 (2008).

21. Fatt, M. P. et al. Restoration of hippocampal neural precursor function by ablation of senescent cells in the aging stem cell niche. Stem Cell Rep. 17, 259–275 (2022).

22. Von Zglinicki, T., Wan, T. & Miwa, S. Senescence in Post-Mitotic Cells: A Driver of Aging? Antioxid. Redox Signal. 34, 308–323 (2021).

23. Dimri, G. P. et al. A biomarker that identifies senescent human cells in culture and in aging skin in vivo. Proc. Natl. Acad. Sci. 92, 9363–9367 (1995).

24. Allard, S., Scardochio, T., Cuello, A. C. & Ribeiro-da-Silva, A. Correlation of cognitive performance and morphological changes in neocortical pyramidal neurons in aging. Neurobiol. Aging 33, 1466–1480 (2012).

25. Kerloch, T., Clavreul, S., Goron, A., Abrous, D. N. & Pacary, E. Dentate Granule Neurons Generated During Perinatal Life Display Distinct Morphological Features Compared With Later-Born Neurons in the Mouse Hippocampus. Cereb. Cortex N. Y. N 1991 29, 3527–3539 (2019).

26. Chen, X. et al. Mass of the postsynaptic density and enumeration of three key molecules. Proc. Natl. Acad. Sci. 102, 11551–11556 (2005).

27. Cameron, H. A., Kaliszewski, C. K. & Greer, C. A. Organization of mitochondria in olfactory bulb granule cell dendritic spines. Synap. N. Y. N 8, 107–118 (1991).

28. Datta, S. & Jaiswal, M. Mitochondrial calcium at the synapse. Mitochondrion 59, 135–153 (2021).

29. Li, Z., Okamoto, K.-I., Hayashi, Y. & Sheng, M. The importance of dendritic mitochondria in the morphogenesis and plasticity of spines and synapses. Cell 119, 873–887 (2004).

30. Masachs, N. et al. The temporal origin of dentate granule neurons dictates their role in spatial memory. Mol. Psychiatry 26, 7130–7140 (2021).

31. Wang, X. & Schwarz, T. L. The Mechanism of Ca2+-Dependent Regulation of Kinesin-Mediated Mitochondrial Motility. Cell 136, 163–174 (2009).

32. Deng, M. et al. Mossy cell synaptic dysfunction causes memory imprecision via miR-128 inhibition of STIM2 in Alzheimer’s disease mouse model. Aging Cell 19, (2020).

33. Li, X. et al. A circuit of mossy cells controls the efficacy of memory retrieval by Gria2I inhibition of Gria2. Cell Rep. 34, 108741 (2021).

34. Li, S. et al. Alzheimer-like tau accumulation in dentate gyrus mossy cells induces spatial cognitive deficits by disrupting multiple memory-related signaling and inhibiting local neural circuit. Aging Cell 21, (2022).

35. Li, Y. et al. Supramammillary nucleus synchronizes with dentate gyrus to regulate spatial memory retrieval through glutamate release. eLife 9, e53129 (2020).

36. Li et al. Hypothalamic modulation of adult hippocampal neurogenesis in mice confers activity-dependent regulation of memory and anxiety-like behavior. Nat. Neurosci. 25, 630–645 (2022).

37. Geinisman, Y., de Toledo-Morrell, L. & Morrell, F. Loss of perforated synapses in the dentate gyrus: morphological substrate of memory deficit in aged rats. Proc. Natl. Acad. Sci. U. S. A. 83, 3027– 3031 (1986).

38. Smith, T. D., Adams, M. M., Gallagher, M., Morrison, J. H. & Rapp, P. R. Circuit-specific alterations in hippocampal synaptophysin immunoreactivity predict spatial learning impairment in aged rats. J. Neurosci. Off. J. Soc. Neurosci. 20, 6587–6593 (2000).

39. Amaral, D. G., Scharfman, H. E. & Lavenex, P. The dentate gyrus: fundamental neuroanatomical organization (dentate gyrus for dummies). Prog. Brain Res. 163, 3–22 (2007).

40. Fischer, W., Gage, F. H. & Björklund, A. Degenerative Changes in Forebrain Cholinergic Nuclei Correlate with Cognitive Impairments in Aged Rats. Eur. J. Neurosci. 1, 34–45 (1989).

41. Lods, M. et al. Adult-born neurons immature during learning are necessary for remote memory reconsolidation in rats. Nat. Commun. 12, 1778 (2021).

42. Lods, M. et al. Chemogenetic stimulation of adult neurogenesis, and not neonatal neurogenesis, is sufficient to improve long-term memory accuracy. Prog. Neurobiol. 219, 102364 (2022).

43. Tronel, S., Lemaire, V., Charrier, V., Montaron, M.-F. & Abrous, D. N. Influence of ontogenetic age on the role of dentate granule neurons. Brain Struct. Funct. 220, 645–661 (2015).

44. Takamori, S. VGLUTs: ‘Exciting’ times for glutamatergic research? Neurosci. Res. 55, 343– 351 (2006).

45. Lagace, D. C. et al. Dynamic Contribution of Nestin-Expressing Stem Cells to Adult Neurogenesis. J. Neurosci. 27, 12623–12629 (2007).

46. Ninkovic, J., Mori, T. & Götz, M. Distinct Modes of Neuron Addition in Adult Mouse Neurogenesis. J. Neurosci. 27, 10906–10911 (2007).

47. Dellu, F., Mayo, W., Vallée, M., Le Moal, M. & Simon, H. Reactivity to novelty during youth as a predictive factor of cognitive impairment in the elderly: a longitudinal study in rats. Brain Res. 653, 51–56 (1994).

48. Dellu, F. et al. Behavioral reactivity to novelty during youth as a predictive factor of stress-induced corticosterone secretion in the elderly--a life-span study in rats. Psychoneuroendocrinology 21, 441–453 (1996).

49. Choi, G. E. et al. BNIP3L/NIX-mediated mitophagy protects against glucocorticoid-induced synapse defects. Nat. Commun. 12, 487 (2021).

50. Choi, G. E. & Han, H. J. Glucocorticoid impairs mitochondrial quality control in neurons. Neurobiol. Dis. 152, 105301 (2021).

51. Murphy, T. & Thuret, S. The systemic milieu as a mediator of dietary influence on stem cell function during ageing. Ageing Res. Rev. 19, 53–64 (2015).

52. Villeda, S. A. et al. The ageing systemic milieu negatively regulates neurogenesis and cognitive function. Nature 477, 90–94 (2011).

53. Smith, L. K. et al. β2-microglobulin is a systemic pro-aging factor that impairs cognitive function and neurogenesis. Nat. Med. 21, 932–937 (2015).

54. Horowitz, A. M. et al. Blood factors transfer beneficial effects of exercise on neurogenesis and cognition to the aged brain. Science 369, 167–173 (2020).

55. Islam, M. R. et al. Exercise hormone irisin is a critical regulator of cognitive function. Nat. Metab. 3, 1058–1070 (2021).

56. Glatigny, M. et al. Autophagy Is Required for Memory Formation and Reverses Age-Related Memory Decline. Curr. Biol. CB 29, 435–448.e8 (2019).

57. Sparta, D. R. et al. Construction of implantable optical fibers for long-term optogenetic manipulation of neural circuits. Nat. Protoc. 7, 12–23 (2011).

58. Drapeau, E. et al. Spatial memory performances of aged rats in the water maze predict levels of hippocampal neurogenesis. Proc. Natl. Acad. Sci. 100, 14385–14390 (2003).

