## Supplementary data for "Long-lived adult-born hippocampal neurons promote successful cognitive aging"

### Supplementary figures S1 to S6

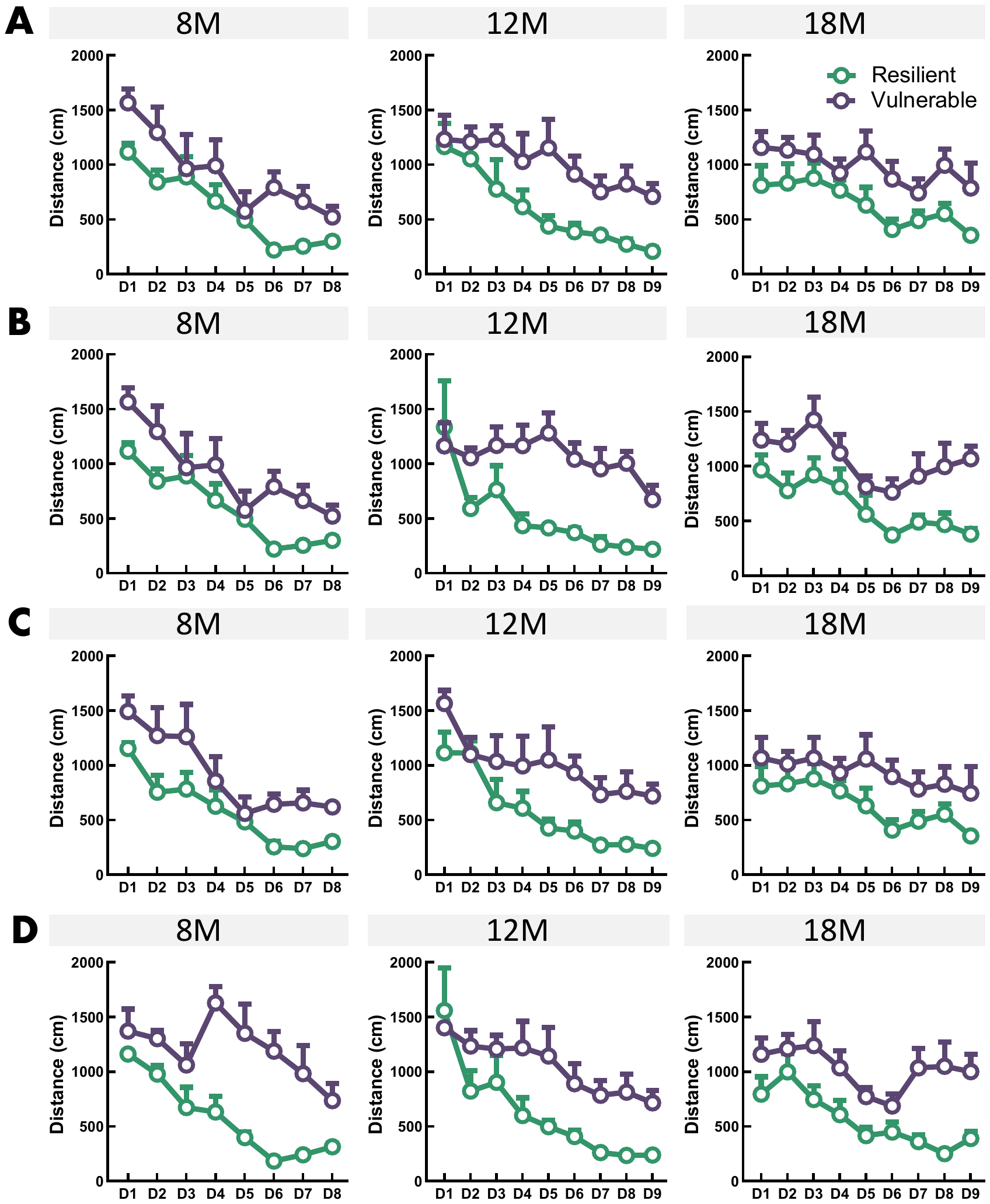
**Figure S1: Resilient animals show better learning abilities than vulnerable animals at each age.** (A) Extremes animals for the two cognitive populations analyzed in the morphology study with M-rv-CAG-GFP. (B) Extremes animals for the two cognitive populations analyzed in the glutamatergic innervation study with M-rv-PSD95-GFP. (C) Extremes animals for the two cognitive populations analyzed in the mitochondrial network study with M-rv-MitoDsRed. (D) Extremes animals for the two cognitive populations analyzed in the survival and senescence study with BrdU and SAβGal.

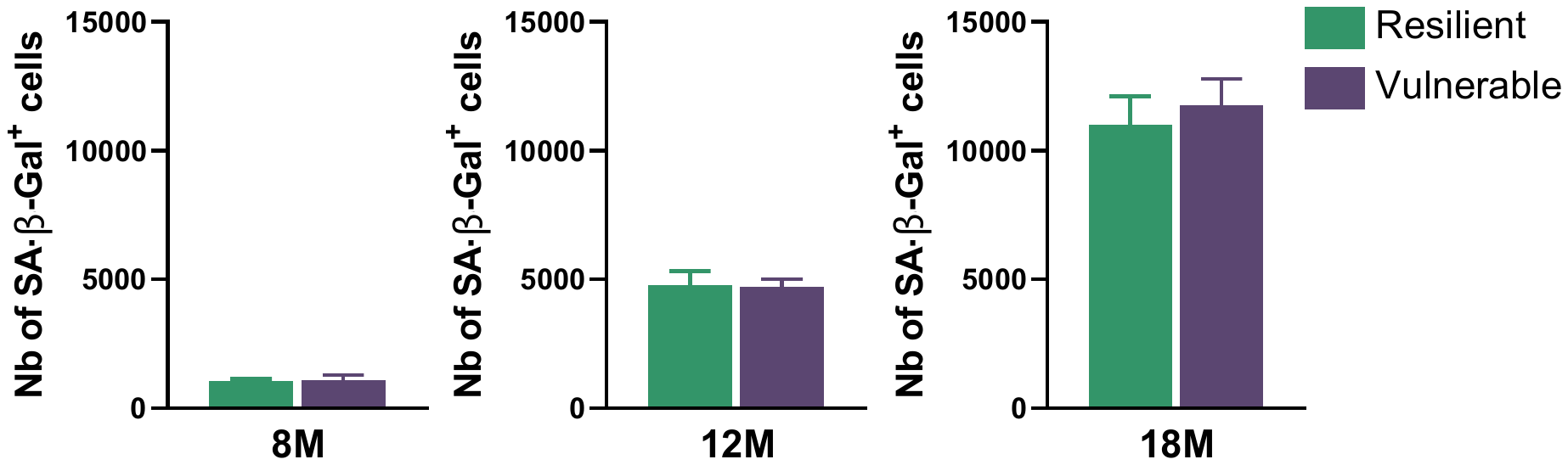
**Figure S2: Cell senescence in the granule cell layer of the dentate gyrus do not account for successful cognitive aging.** Resilient and vulnerable animals show similar number of senescent cells in the granule cell layer at 8-month-old (unpaired *t* test: *t_8_* = 0.28, P > 0.05), 12-month-old (unpaired *t* test: *t_8_* = 0.14, P > 0.05) and 18-month-old (unpaired *t* test: *t_8_* = 0.51, P > 0.05). Data are presented as mean ± S.E.M. from 5 extreme animals per group. Statistical significance *P ≤ 0.05, **P < 0.01, ***P < 0.001.

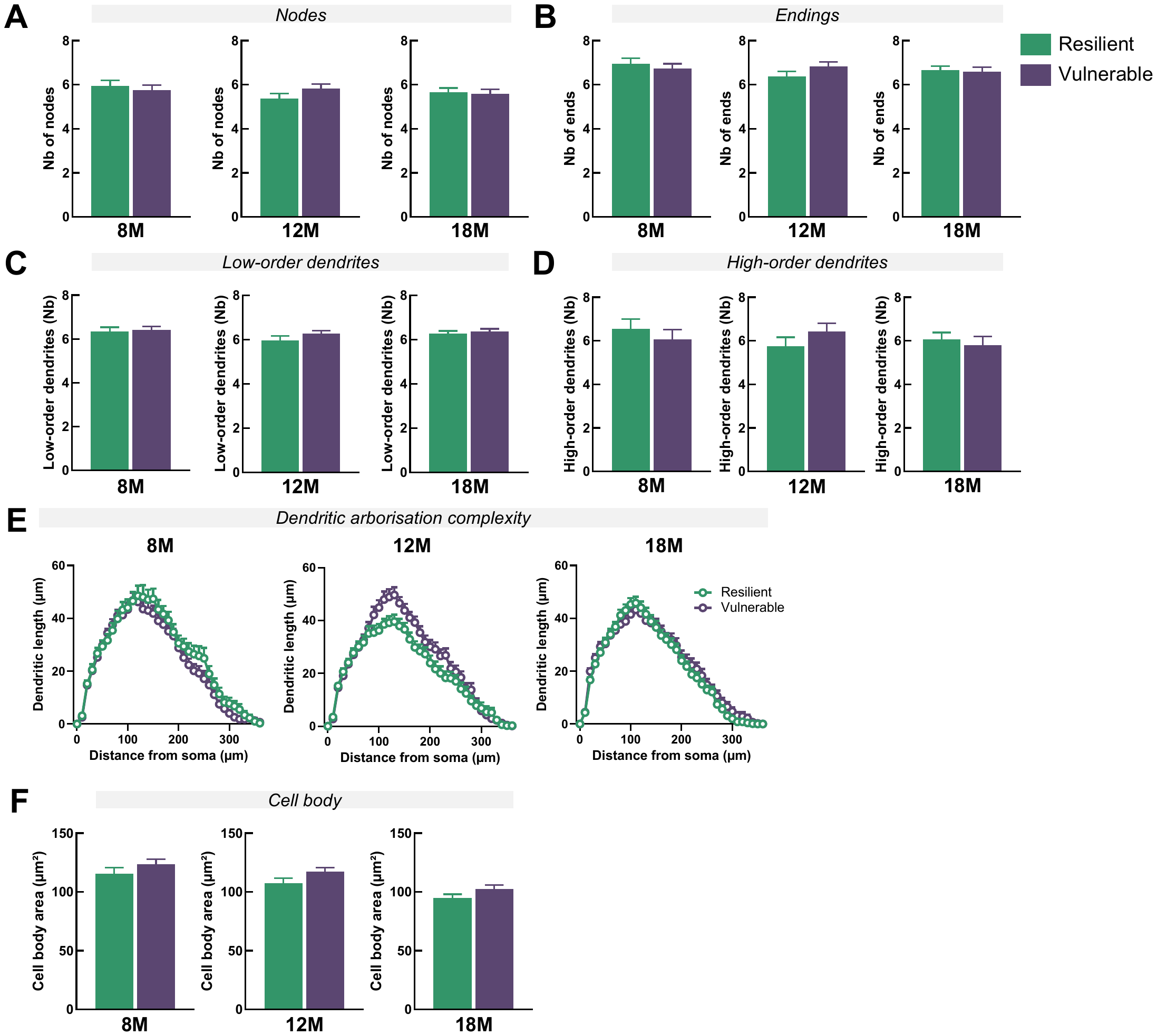
**Figure S3: The gross morphological architecture of ABNs is independent of the cognitive status of the rats** (A) ABNs in the two cognitive populations show similar number of nodes at 8-month-old (unpaired *t* test: *t_90_* = 0.55, P > 0.05), 12-month-old (unpaired *t* test: *t_102_* = 1.59, P > 0.05) and 18-month-old (unpaired *t* test: *t_111_* = 0.31, P > 0.05). (B) ABNs have similar number of ends in resilient and vulnerable animals at 8-month-old (unpaired *t* test: *t_90_* = 0.64, P > 0.05), 12-month-old (unpaired *t* test: *t_102_* = 1.53, P > 0.05) and 18-month-old (unpaired *t* test: *t_111_* = 0.31, P > 0.05). (C) ABNs in the two cognitive populations show similar low-order dendrites number at 8-month-old (unpaired *t* test: *t_90_* = 0.28, P > 0.05), 12-month-old (unpaired *t* test: *t_102_* = 1.33, P > 0.05) and 18-month-old (unpaired *t* test: *t_111_* = 0.48, P > 0.05). (D) ABNs show similar high-order dendrites number at 8-month-old (unpaired *t* test: *t_90_* = 0.72, P > 0.05), 12-month-old (unpaired *t* test: *t_102_* = 1.22, P > 0.05) and 18-month-old (unpaired *t* test: *t_111_* = 0.02, P > 0.05). (E) ABNs have similar dendritic complexity between the two cognitive populations at 8-month-old (RM-two-way ANOVA, F_36,3240_=0.68, P > 0.05) and 18-month-old (RM-two-way ANOVA, F_36,3996_=0.82, P > 0.05). At 12-month-old, a small difference could be detected between the resilient and vulnerable populations (RM-two-way ANOVA, F_36,3708_=2.59, P < 0.001). (F) ABNs in the two cognitive populations show similar cell body area at 8-month-old (unpaired *t* test: *t_90_* = 1.22, P > 0.05), 12-month-old (unpaired *t* test: *t_102_* = 1.84, P > 0.05) and 18-month-old (unpaired *t* test: *t_111_* = 1.66, P > 0.05). Data are presented as mean ± S.E.M. from 5 extreme animal per groups (a min of 4 neurons were traced per animal, with 8M-Res = 37 neurons and 8M-Vul = 55 neurons; 12M-Res = 48 neurons and 12M-Vul = 58 neurons; 18M-Res = 63 neurons and 18M-Vul = 50 neurons. Statistical significance *P ≤ 0.05, **P < 0.01, ***P < 0.001.

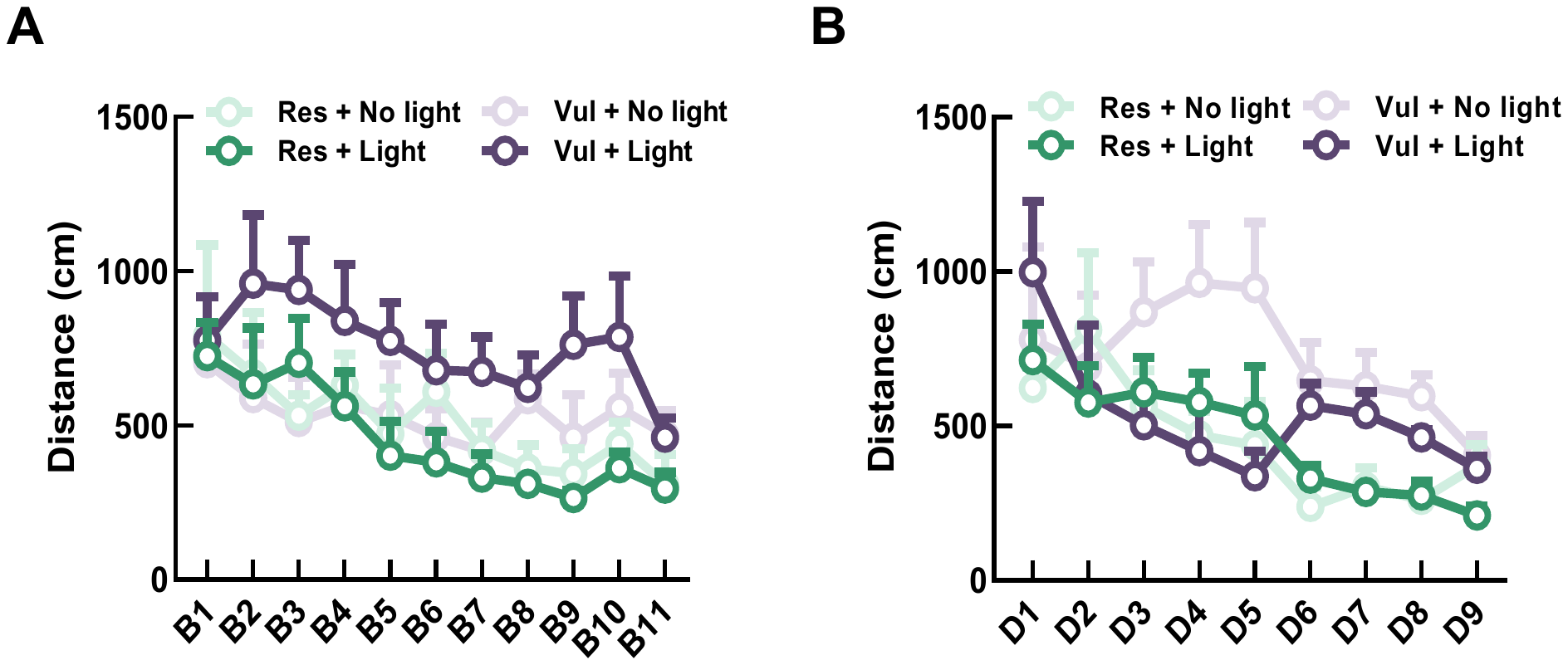
**Figure S4: Similar levels of ABNs stimulation between resilient and vulnerable animals.** (A) Similar percentage of ABNs stimulation between resilient and vulnerable animals at 20-month-old. (B)
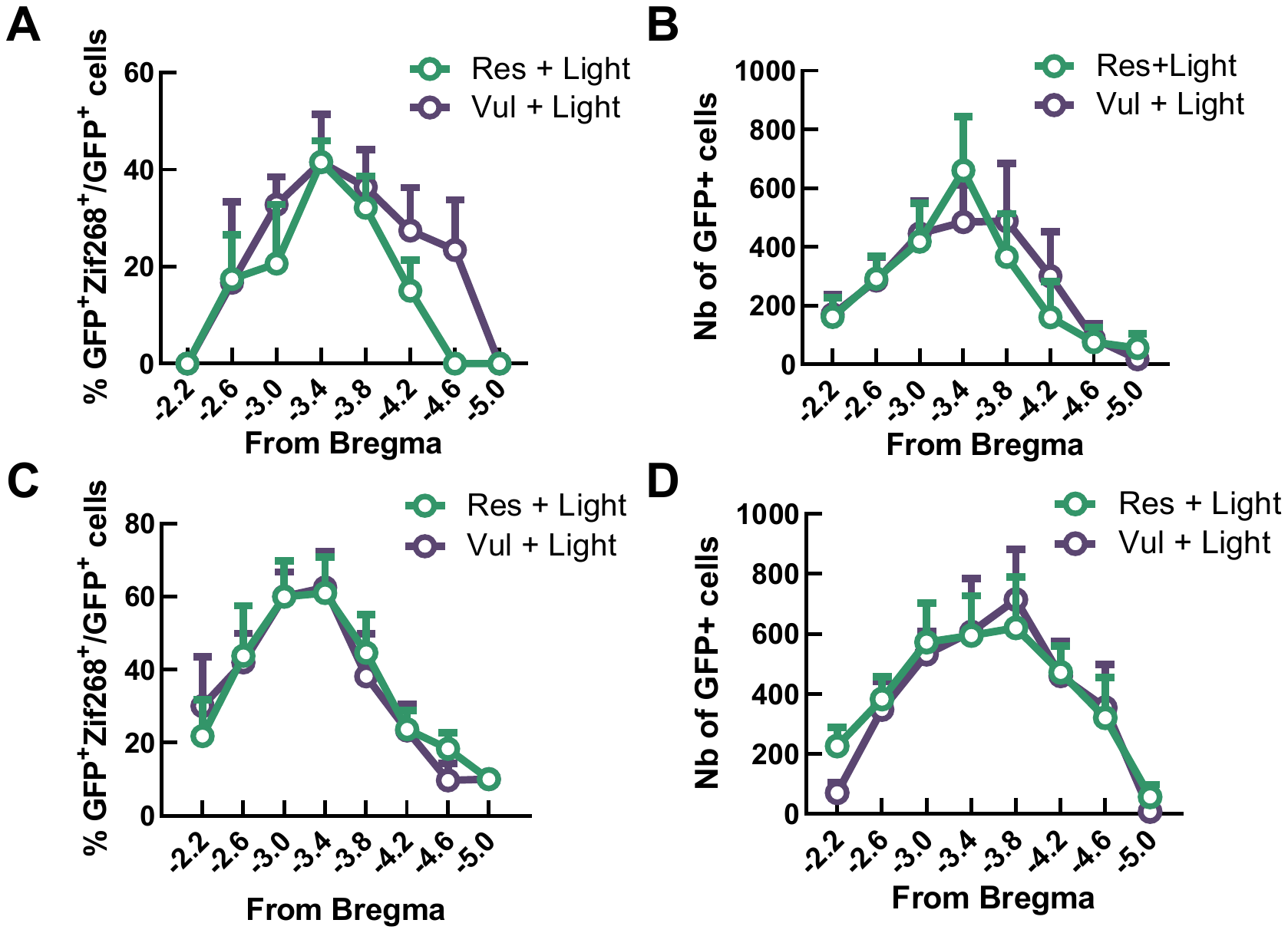
Similar number of GFP labelled ABNs between stimulated resilient and vulnerable animals at 20-month-old. (C) Similar percentage of ABNs stimulation between resilient and vulnerable animals at 12-month-old. (D) Similar number of GFP labelled ABNs between stimulated resilient and vulnerable animals at 12-month-old. Data are presented as mean ± S.E.M for 20M-Res-Light: 5 animals, 20M-Vul-Light: 4 animals,12M-Res-Light: 6 animals, 12M-Vul-Light: 5 animals. For the percentage of activation, data expressed as (Nb of GFP^+^-Zif268^+^ cells)/(Nb of GFP^+^- Zif268^−^ cells + Nb of GFP^+^- Zif268^+^ cells) × 100.

**Figure S5: Stimulation of ABNs during the learning phase has no effect on the performances.** (A) Learning performances of each group over several days of training at 20-month-old. Days of learning are presented in blocks; one block corresponds to two days of learning (B) Learning performances of each group over several days of training at 12-month-old. Data are presented as mean ± S.E.M for: 5 animals, 20M-Res-Light: 5 animals; 20M-Vul-Light: 4 animals, 20M-Res-NoLight: 4 animals, 20M-Vul-NoLight: 4 animals, 12M-Res-Light: 6 animals, 12M-Vul-Light: 5 animals, 12M-Res-NoLight: 5 animals and 12M-Vul-NoLight: 4 animals.

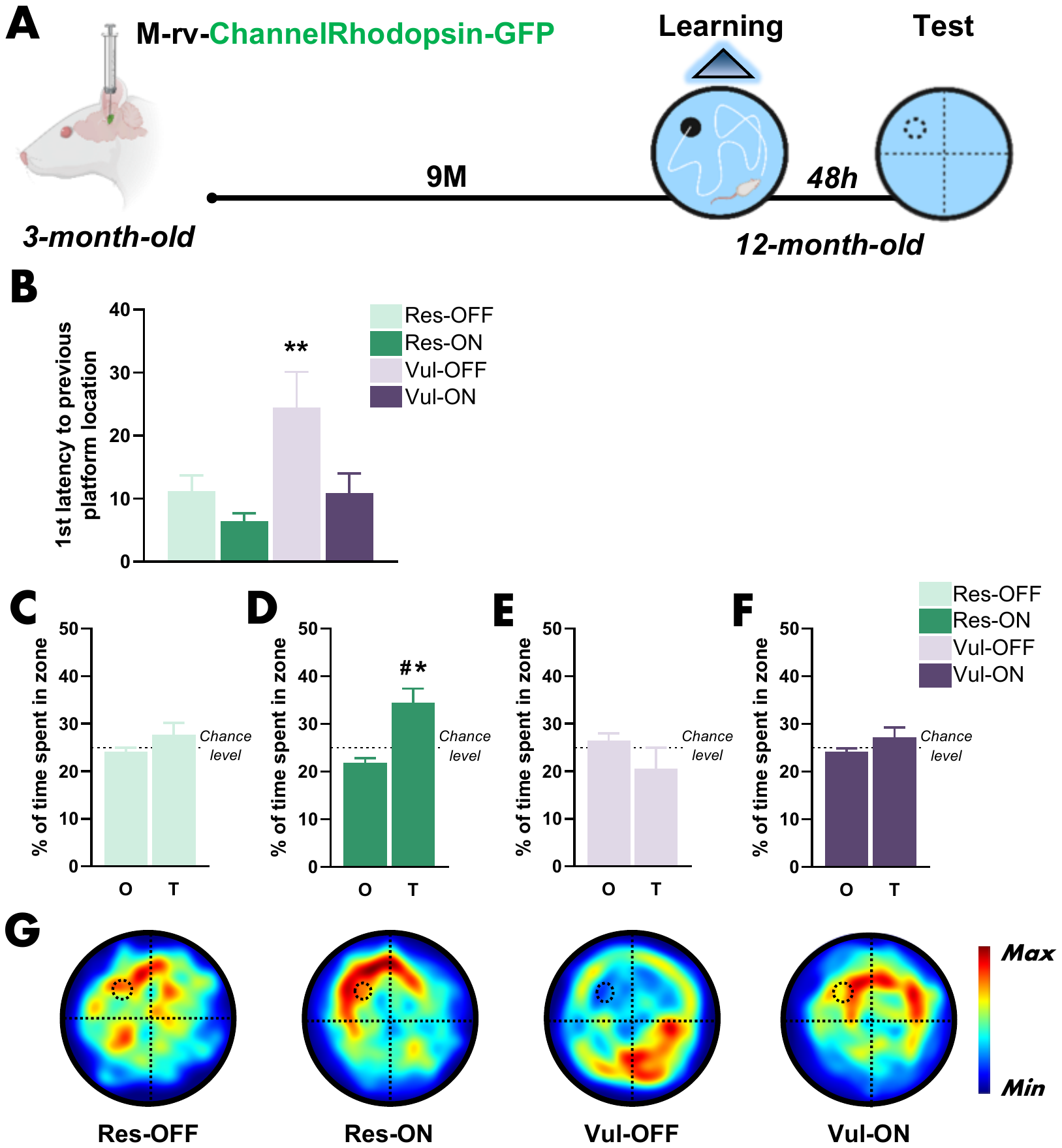

**Figure S6: Stimulation of ABNs during learning restores memory retrieval in middle-aged vulnerable animals.** (A) Schematic diagram of the experimental design. (B) Stimulation of ABNs in middle-age stimulated vulnerable (Vul-ON) animals promoted the formation of a precise memory trace similar to resilient animals (Res-OFF and Res-ON) (One-way-ANOVA, F_(3, 16)_ = 5,94, P < 0.01). (C) Middle-age non-stimulated resilient (Res-OFF) did not form a stable memory trace (paired *t* test: *t_4_* = 1.08, P > 0.05). (D) Stimulation of ABNs in middle-age stimulated resilient (Res-ON) animals promoted the formation of a strong memory trace (paired *t* test: *t_5_* = 3.23, P ≤ 0.05 and one sample *t* test against ^#^chance level *t*_5_ = 3,23, P < 0,001). (E) Middle-age non-stimulated vulnerable (Vul-OFF) animals did not form a stable memory trace (paired *t* test: *t_3_* = 1.01, P > 0.05). (F) Stimulation of ABNs did not restore the formation of a stable memory trace in Vul-ON animals (paired *t* test: *t_4_* = 1.15, P > 0.05). (G) Heatmaps depicting the individuals search location and occupancy during the memory test. Data are presented as mean ± S.E.M. Statistical significance *P ≤ 0.05, **P < 0.01, ***P < 0.001.

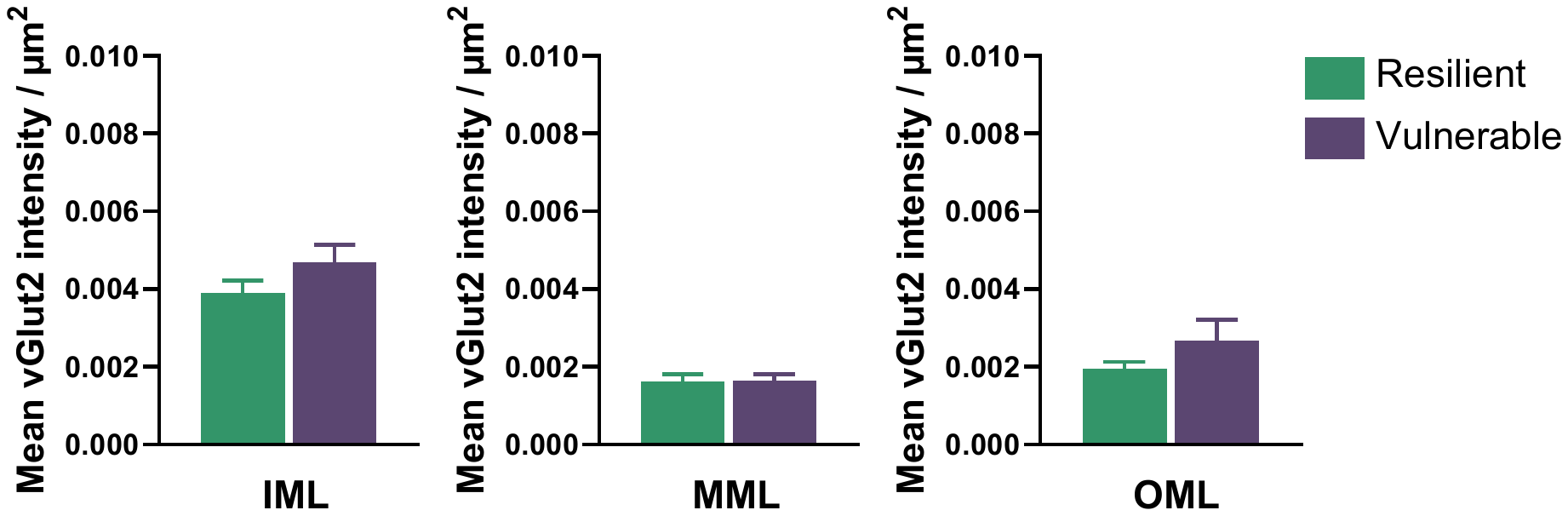
**Figure S7: Similar glutamatergic innervation in the molecular layer of the dentate gyrus between resilient and vulnerable individuals at old age.** Resilient and vulnerable animals show similar mean intensity for vGLUT2 labelling at 18-month-old in the IML (unpaired *t* test_25_: *t* = 1.38 P > 0.05), MML (unpaired *t* test: *t*_25_ = 0.09, P > 0.05) and OML (unpaired *t* test: *t_25_* = 1.12, P > 0.05). Data are presented as mean ± S.E.M. from 5 extreme animals per group. Statistical significance *P ≤ 0.05, **P < 0.01, ***P < 0.001.

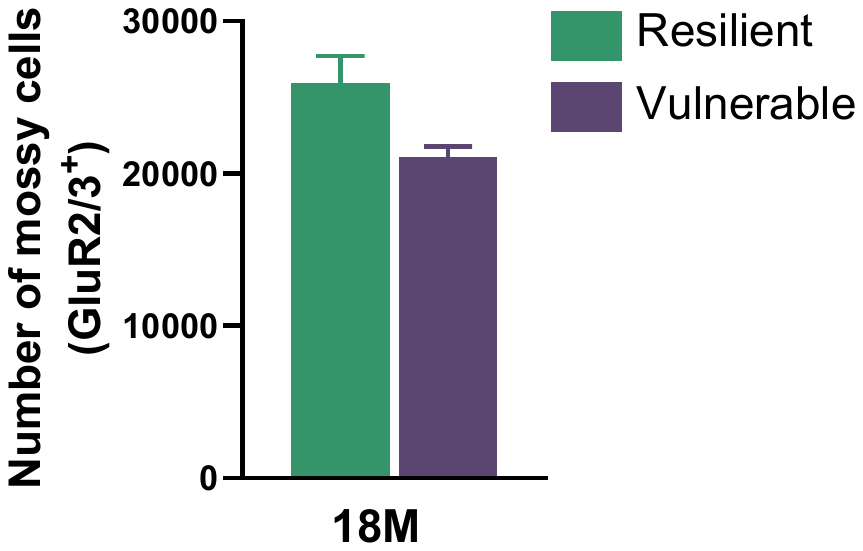
**Figure S8: Similar number of mossy cells in the hilus of the dentate gyrus between resilient and vulnerable individuals at old age.** Resilient and vulnerable animals show similar number of mossy cells expressing GluR2/3 at 18-month-old (unpaired *t* test: *t_4_* = 2.57, P > 0.05). Data are presented as mean ± S.E.M. from 3 extreme animals per group. Statistical significance *P ≤ 0.05, **P < 0.01, ***P < 0.001.
